# A sequential two-step priming scheme reproduces diversity in synaptic strength and short-term plasticity

**DOI:** 10.1101/2022.05.08.491082

**Authors:** Kun-Han Lin, Holger Taschenberger, Erwin Neher

## Abstract

Glutamatergic synapses display variable strength and diverse short-term plasticity (STP), even for a given type of connection. Using non-negative tensor factorization (NTF) and conventional state modelling, we demonstrate that a kinetic scheme consisting of two sequential and reversible steps of release-machinery assembly and a final step of synaptic vesicle (SV) fusion reproduces STP and its diversity among synapses. Analyzing transmission at calyx of Held synapses reveals that differences in synaptic strength and STP are not primarily caused by variable fusion probability (*p_fusion_*) but determined by the fraction of docked synaptic vesicles equipped with a mature release machinery. Our simulations show, that traditional quantal analysis methods do not necessarily report *p_fusion_* of SVs with a mature release machinery but reflect both *p_fusion_* and the distribution between mature and immature priming states at rest. Thus, the approach holds promise for a better mechanistic dissection of the roles of presynaptic proteins in the sequence of SV docking, two-step priming and fusion and suggests a mechanism for activity-induced redistribution of synaptic efficacy.

## Introduction

Chemical synapses change their strength during repetitive use in a synapse type-specific and activity-dependent manner. Such modifications can occur on several time scales and define dynamic properties of synaptic networks (1–3). Elucidating the biophysical mechanisms of synaptic plasticity is essential to understand information processing in circuits (4–6). Kinetic schemes of synaptic transmission and plasticity also provide a theoretical framework to mechanistically and quantitatively interpret functional synaptic deficits due to molecular perturbations of the release machinery, either experimentally induced or arising as a consequence of genetically determined synaptopathies (7).

Short-term changes of synaptic strength such as paired-pulse facilitation (PPF) and short-term depression (STD) have been ascribed to changes in the release probability of fusion-competent synaptic vesicles (SVs) and/or changes in the occupancy of a fixed number of presynaptic release sites (8, 9). Some basic features of short-term plasticity (STP) are captured by a simple scheme postulating one kind of release site to which SVs are recruited, possibly in a Ca^2+-depen^dent manner, before being able to fuse upon action potential (AP) arrival (10, 11).

However, a number of experimental observations, including multiple kinetic components of STD (12) and its recovery (13, 14) and diverse STP even among synapses of a given type (15–18) are not easily accounted for by such a simple model restricted to a single functionally homogenous pool of fusion-competent SVs.

To more faithfully reproduce the multifaceted features of STP, different multiple-state and/or multiple-site schemes of transmitter release have been proposed (19–26), which fall into two principal categories: (*i*) parallel schemes in which more than one kind of SVs can bind to one or more kinds of release sites, or (*ii*) sequential schemes in which SVs migrate between different kinds of release sites, or else undergo changes of maturation, reflected by distinct SV states, while being docked to a given site.

Motivated by converging evidence from molecular biology (27–31) electrophysiology (32–35), live-cell imaging (36, 37) and electron microscopy (EM) on samples prepared by cryofixation (38, 39) emphasizing reversibility and multistep nature of the SV priming process, we explore here whether a recently proposed single-site multiple-state scheme of priming and fusion (40) can reproduce the variable synaptic strength and diverse STP observed at the calyx of Held, a large glutamatergic nerve terminal in the mammalian brainstem. The proposed kinetic scheme in its basic form (Fig. 1A,B) assumes that SVs reversibly dock to a single type of release site and undergo two sequential priming steps to become fusion competent. In view of the ultrastructural evidence for distinct docking states (38, 41–44), we refer to the two states and to the SVs residing in those states as loosely (LS) or tightly (TS) docked and SVLS or SVTS, respectively.

**Figure 1.**
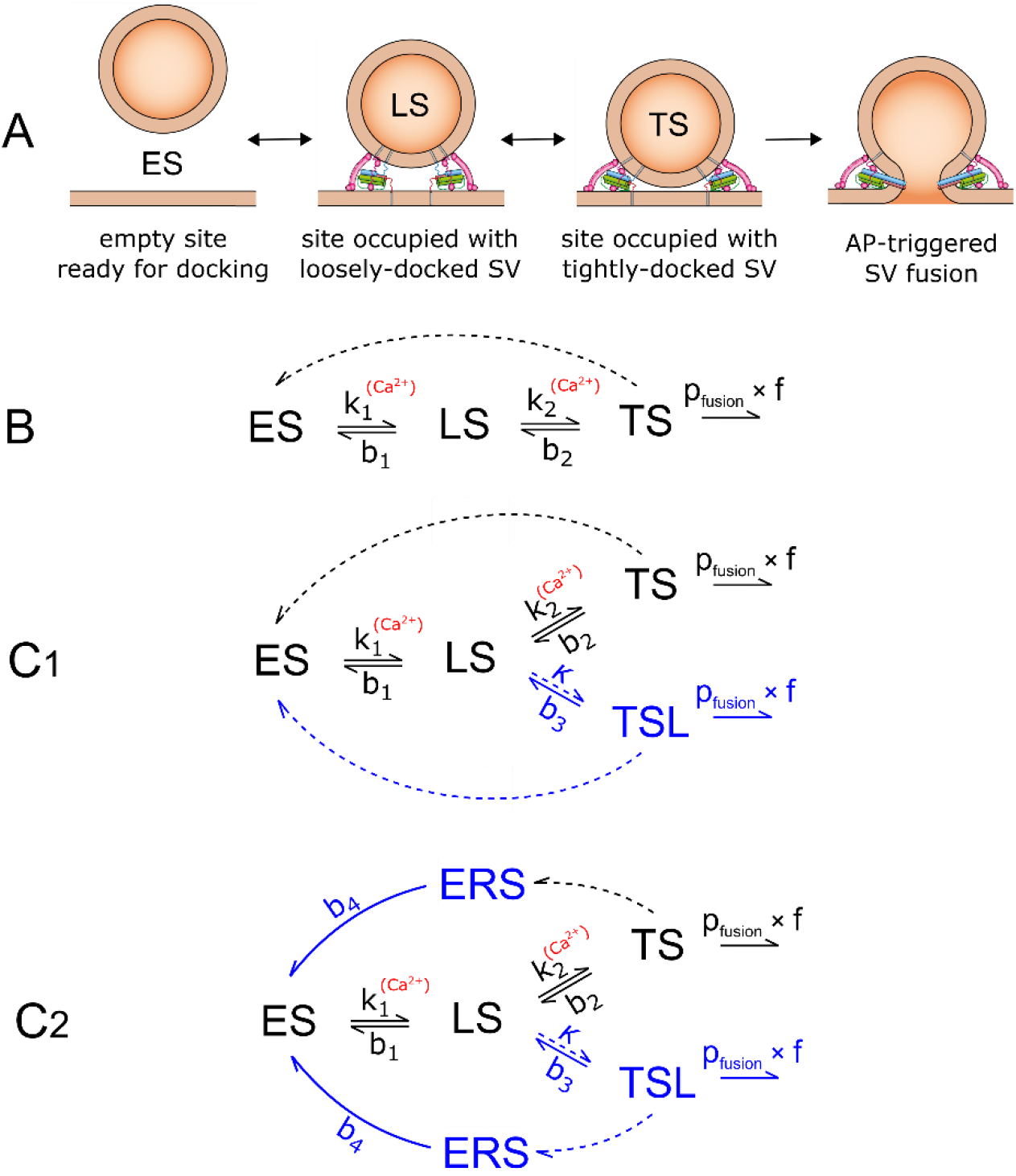
Diagram of vesicle states and kinetic schemes for the numerical simulation of STP. **A:** Basic sequential model for SV priming and fusion. SVs dock to an empty release site (ES) and undergo two priming steps to sequentially transition to the states LS (loosely docked) and TS (tightly docked). Only SVs in state TS are fusion-competent. **B:** Kinetic scheme of state transitions for the basic model shown in **A**. A simple three-state scheme is adequate for reproducing experimental results using *f_stim_* = 1–20 Hz. **C:** An extended reaction scheme with an additional labile tightly docked state (TSL) is required for reproducing experimental results for *f_stim_* ≥50 Hz. SV_TSs_ and SVTSLs fuse with the same *p_fusion_* in response to an AP. The two states TS and TSL differ with respect to their stability. While TS has a lifetime in the range of 3–4 s, TSL relaxes back to LS within ∼100 ms. Vacated release sites can either instantaneously return to ES and thereby be immediately available for SV docking (**C1**), or else reside for some time in a refractory empty state (ERS) (**C2**). State transitions in **B** and **C** represented by dashed lines indicate instantaneous transitions, while those represented by solid lines occur with rate constants as shown (see also Table 1). Elements shown in blue in **C** extend the kinetic scheme illustrated in **B**.

**Table 1.**
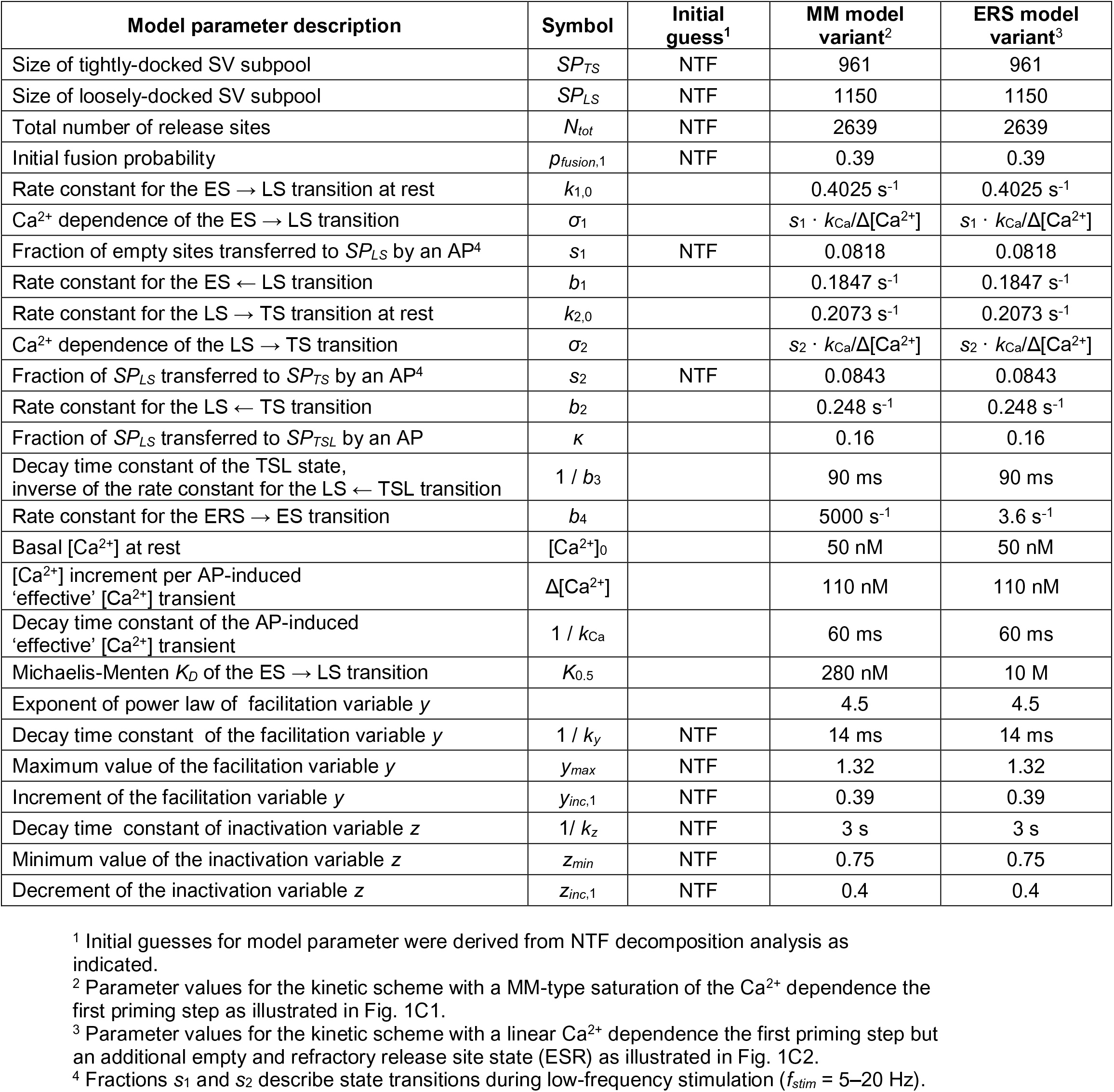
Model parameter descriptions and parameter values for the sequential two-step priming and fusion scheme.

| Model parameter description | Symbol | Initial guess<sup>1</sup> | MM model variant<sup>2</sup> | ERS model variant<sup>3</sup> |
| --- | --- | --- | --- | --- |
| Size of tightly-docked SV subpool | $SP_{TS}$ | NTF | 961 | 961 |
| Size of loosely-docked SV subpool | $SP_{LS}$ | NTF | 1150 | 1150 |
| Total number of release sites | $N_{tot}$ | NTF | 2639 | 2639 |
| Initial fusion probability | $p_{fusion,1}$ | NTF | 0.39 | 0.39 |
| Rate constant for the ES → LS transition at rest | $k_{1,0}$ | | 0.4025 s<sup>-1</sup> | 0.4025 s<sup>-1</sup> |
| Ca<sup>2+</sup> dependence of the ES → LS transition | $\sigma_1$ | | $s_1 \cdot k_{Ca}/\Delta[Ca^{2+}]$ | $s_1 \cdot k_{Ca}/\Delta[Ca^{2+}]$ |
| Fraction of empty sites transferred to $SP_{LS}$ by an AP<sup>4</sup> | $s_1$ | NTF | 0.0818 | 0.0818 |
| Rate constant for the ES ← LS transition | $b_1$ | | 0.1847 s<sup>-1</sup> | 0.1847 s<sup>-1</sup> |
| Rate constant for the LS → TS transition at rest | $k_{2,0}$ | | 0.2073 s<sup>-1</sup> | 0.2073 s<sup>-1</sup> |
| Ca<sup>2+</sup> dependence of the LS → TS transition | $\sigma_2$ | | $s_2 \cdot k_{Ca}/\Delta[Ca^{2+}]$ | $s_2 \cdot k_{Ca}/\Delta[Ca^{2+}]$ |
| Fraction of $SP_{LS}$ transferred to $SP_{TS}$ by an AP<sup>4</sup> | $s_2$ | NTF | 0.0843 | 0.0843 |
| Rate constant for the LS ← TS transition | $b_2$ | | 0.248 s<sup>-1</sup> | 0.248 s<sup>-1</sup> |
| Fraction of $SP_{LS}$ transferred to $SP_{TSL}$ by an AP | $\kappa$ | | 0.16 | 0.16 |
| Decay time constant of the TSL state, inverse of the rate constant for the LS ← TSL transition | $1 / b_3$ | | 90 ms | 90 ms |
| Rate constant for the ERS → ES transition | $b_4$ | | 5000 s<sup>-1</sup> | 3.6 s<sup>-1</sup> |
| Basal [Ca<sup>2+</sup>] at rest | $[Ca^{2+}]_0$ | | 50 nM | 50 nM |
| [Ca<sup>2+</sup>] increment per AP-induced 'effective' [Ca<sup>2+</sup>] transient | $\Delta[Ca^{2+}]$ | | 110 nM | 110 nM |
| Decay time constant of the AP-induced 'effective' [Ca<sup>2+</sup>] transient | $1 / k_{Ca}$ | | 60 ms | 60 ms |
| Michaelis-Menten $K_D$ of the ES → LS transition | $K_{0.5}$ | | 280 nM | 10 M |
| Exponent of power law of facilitation variable $y$ | | | 4.5 | 4.5 |
| Decay time constant of the facilitation variable $y$ | $1 / k_y$ | NTF | 14 ms | 14 ms |
| Maximum value of the facilitation variable $y$ | $y_{max}$ | NTF | 1.32 | 1.32 |
| Increment of the facilitation variable $y$ | $y_{inc,1}$ | NTF | 0.39 | 0.39 |
| Decay time constant of inactivation variable $z$ | $1 / k_z$ | NTF | 3 s | 3 s |
| Minimum value of the inactivation variable $z$ | $z_{min}$ | NTF | 0.75 | 0.75 |
| Decrement of the inactivation variable $z$ | $z_{inc,1}$ | NTF | 0.4 | 0.4 |

<sup>1</sup> Initial guesses for model parameter were derived from NTF decomposition analysis as indicated.
<sup>2</sup> Parameter values for the kinetic scheme with a MM-type saturation of the Ca<sup>2+</sup> dependence the first priming step as illustrated in Fig. 1C1.
<sup>3</sup> Parameter values for the kinetic scheme with a linear Ca<sup>2+</sup> dependence the first priming step but an additional empty and refractory release site state (ERS) as illustrated in Fig. 1C2.
<sup>4</sup> Fractions $s_1$ and $s_2$ describe state transitions during low-frequency stimulation ( $f_{stim} = 5\text{--}20$ Hz).

We analyzed eEPSC trains elicited by a wide range of presynaptic firing frequencies using a combined electrophysiological and modelling approach which was aided by non-negative tensor factorization (NTF) (45). Our analysis leads to four principal conclusions: (1) The experimental data are well compatible with a reversible SV priming process leaving ∼20% of the release sites empty at rest, while the remaining 80% are occupied by either SVLSs or SV_TSs_, which constitute SV subpools *SP_LS_* and *SP_TS_*, respectively. Both subpools equilibrate dynamically during presynaptic activity. (2) Different initial strength and diverse STP among calyx synapses is primarily due to variable *SP_LS_* and *SP_TS_* sizes at rest. Functional diversity across all synapses is consistent with relatively uniform *p_fusion_* despite the large variability in initial strength. (3) Fusion-competent docked and primed SV_TSs_ have a high *p_fusion_* of ∼0.4, consistent with the experimentally observed rapidly progressing STD during high-frequency stimulation. (4) Depending on presynaptic discharge frequency, release occurs with different characteristics: (*i*) For low frequencies, consecutive release episodes hardly influence each other, except for changes in the occupancy of subpools: Each AP triggers the fusion of a fraction of SV_TSs_ and causes a forward transition of a constant fraction of SVs to the respective downstream SV subpool, irrespective of the interval between consecutive APs. (*ii*) At high frequencies, additional kinetic features become apparent such as an increase in *p_fusion_* during trains, a speed-up of the priming process, and a frequency-dependent decline of SV subpool occupancy and steady-state release rate. The first two features contribute to PPF, while the third one causes STD. For modelling these high-frequency features, the kinetic scheme has to be extended (Fig. 1C).

Our numerical simulations, which mostly use experimentally determined or NTF- constrained model parameters, faithfully reproduce STP at calyx synapses over a wide range of presynaptic activity levels and therefore provide a valuable framework for mechanistic and quantitative interpretation of experimentally induced STP alterations. By emphasizing the multistep nature and reversibility of the SV priming process, the sequential two-step priming scheme suggests a novel mechanism for creating functional diversity among synapses and for an activity-induced redistribution of synaptic efficacy during AP trains.

## Results

In order to validate the two-step priming scheme, we chose the following three-step approach: (1) We acquired eEPSCs evoked by regular stimulus trains (0.5–200 Hz) from an ensemble of 35 post-hearing onset rat calyx of Held synapses. Additionally, 100 and 200 Hz eEPSC trains preceded by either 2 or 4 stimuli at 10 Hz were recorded. (2) Next, eEPSC train data were subjected to NTF analysis with the aim of deriving suitable initial guesses for various model parameters. (3) Parameters of the release model were then initialized with values derived either from NTF analysis or from analytical expressions regarding model predictions for e.g. *PPR* and steady-state release at low-frequency stimulation (eqns. 22–24) and subsequently optimized by trial-and-error variation to closely reproduce the respective average eEPSC train response for each stimulation frequency (*f_stim_*).

### Experimental eEPSC train data: Mean time courses and variability among calyx synapses

Figure 2A,B illustrates eEPSCs and time courses of their mean quantal content in response to stimulus trains consisting of 15 (0.5, 1 and 2 Hz) or 40 APs (5, 10, 20, 50, 100 and 200 Hz). For a given synapse, the quantal content estimates for the initial eEPSCs (*m*1) were similar across all *f_stim_*. However they varied nearly 10fold between synapses (77–739 SVs, CV = 0.42, Fig. 2D). The average over all 35 mean *m*_1_ values was 377 ± 27 SV (Fig. 2B). During trains, the mean *mj* decreased monotonically towards a depressed steady state for all but the highest *f_stim_*. During 200 Hz stimulation, however, net facilitation was observed for the average response i.e. the paired-pulse-ratio (*PPR*_200 Hz_ = *m*_2_ / *m*_1_) was on average >1 (Fig. S1A). Plotting *PPR*_200_ Hz as a function of *m*_1_ for individual synapses revealed large diversity among synapses and a negative correlation, i.e. synapses with a large *m*_1_ tended to show various degrees of paired-pulse depression while synapses with smaller *m*_1_ often exhibited paired-pulse facilitation (PPF) (Fig. 2D). Such correlation is usually interpreted as an indication of heterogeneous initial *p_fusion_* (*p_fusion_*_,1_), because strong depression in synapses with high *p_fusion_*,_1_ would occlude PPF (46). The two-step priming scheme proposed here (Fig. 1A,B) allows an alternative view: It provides an analytical expression for calculating *p_fusion_*,_1_ of individual synapses from *PPR* and STD during 10 Hz eEPSC trains (eqn. 31). Plotting *p_fusion_*,_1_ estimates versus the respective *m*_1_ values for all synapses revealed only weak positive correlation (Fig. 2E). The mean *p_fusion_*,_1_ amounted to 0.43 with a CV of 0.20. The latter was substantially smaller than the CV of individual *m*_1_ values, indicating that variability of *p_fusion_*,_1_ is not the principal cause for heterogeneous synaptic strength. The number of fusion-competent SVs at rest (*SP_TS_*_,0_) is readily calculated for each synapse as the ratio *m*_1_ / *p_fusion_*_,1_.

**Figure 2.**
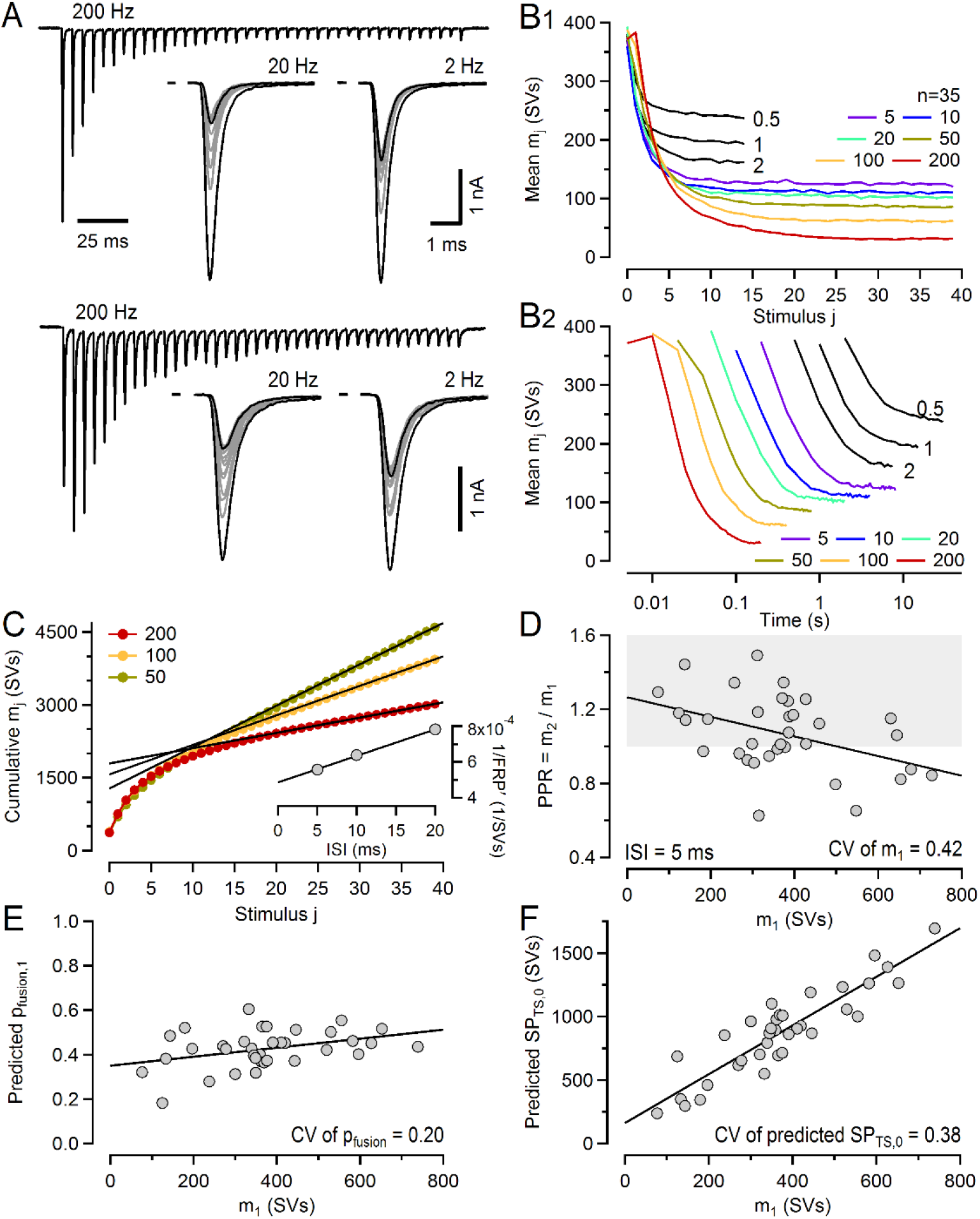
STP in response to 0.5–200 Hz stimulus trains in post-hearing-onset calyx of Held synapses. **A:** Sample eEPSCs obtained from a strongly depressing (top) and a facilitating (bottom) synapse in response to 200 Hz (left), 20 Hz (middle) and 2 Hz (right) stimulation. Only the initial 15 eEPSCs are superimposed for the 2 and 20 Hz eEPSC trains. Each trace represents an average of three repetitions. **B:** Mean quantal content (*mj*) plotted against stimulus index *j* (**B1**) or time (**B2**) for each eEPSC. Trains consisted of only 15 stimuli for the lowest three frequencies. The timing of eEPSC1 was offset by one ISI in **B2** for clarity. Note logarithmic time axis in **B2**. **C:** Estimating the mean *FRP*’ from eEPSC trains evoked by 50, 100 and 200 Hz stimulation. The relationship between the three 1/*FRP*’ values and their respective ISIs was extrapolated to infinite *f_stim_* (ISI = 0 ms) to obtain a mean *FRP* value which is corrected for incomplete pool depletion (inset). **D:** *PPR*s (*m*_2_ / *m*1) for 200 Hz eEPSC trains negatively correlate with initial quantal content (*m*1), which varies approximately tenfold among calyx synapses (73–728 SVs). The gray shaded region indicates *PPR* >1. **E,F.** Predictions for *p_fusion_* (**E**) and *SP*TS,0 (**F**) for individual synapses obtained from their respective 10 Hz *PPR* and *Dm* values according to eqn. 31.

Plotting individual *SPTS,*0 estimates as a function of the respective *m*_1_ values revealed a strong linear correlation with an expected slope of ∼1/0.43 (Fig. 2F). The mean *SPTS,*0 estimate amounted to 880 ± 57 SVs with a CV of 0.38, which is close to the CV of *m*_1_ values. Thus, a large heterogeneity among synapses with respect to their *SP_TS_* size at rest can explain strong variability in *m*1.

### NTF analysis provides pfusion estimates and yields constraints for SV subpool sizes and priming kinetics

We next sought to derive suitable initial guesses for model parameters of the sequential SV priming and fusion scheme (Fig. 1) from NTF analysis of eEPSC trains. NTF decomposes complex data sets into a linear combination of a small number of components (47) and was recently adapted for the analysis of eEPSC trains (45). NTF assumes that time courses of eEPSC peak values during trains are superpositions of contributions by two or more types of signal sources and that their differential relative contributions account for STP diversity among synapses. In the context of the model (Fig. 1), the sources correspond to the release contributions by SVs residing in certain states at the onset of stimulation. NTF analysis returns the time courses of the individual contributions, called basefunctions (BFs). BFs are the same for all synapses. They are normalized to a cumulative sum of 1, such that the product of a BF and the corresponding eEPSC train quantal content (*M*) represents the time-resolved quantal release of that component (Fig. 3B, Fig. S2A, S2B). Two-component NTF fits provide basefunctions for release contributed by pre-existing SV_TSs_ (BF_TS_; Fig. 3A), and for the combined remaining release originating from SVs, which had been either loosely docked at stimulation onset or were newly recruited during the stimulus train (BF_LS,RS_; Fig. 3B). The latter release component can be decomposed by subsequent three-component NTF analysis (Fig. 3C,Fig. S2A,B).

**Figure 3.**
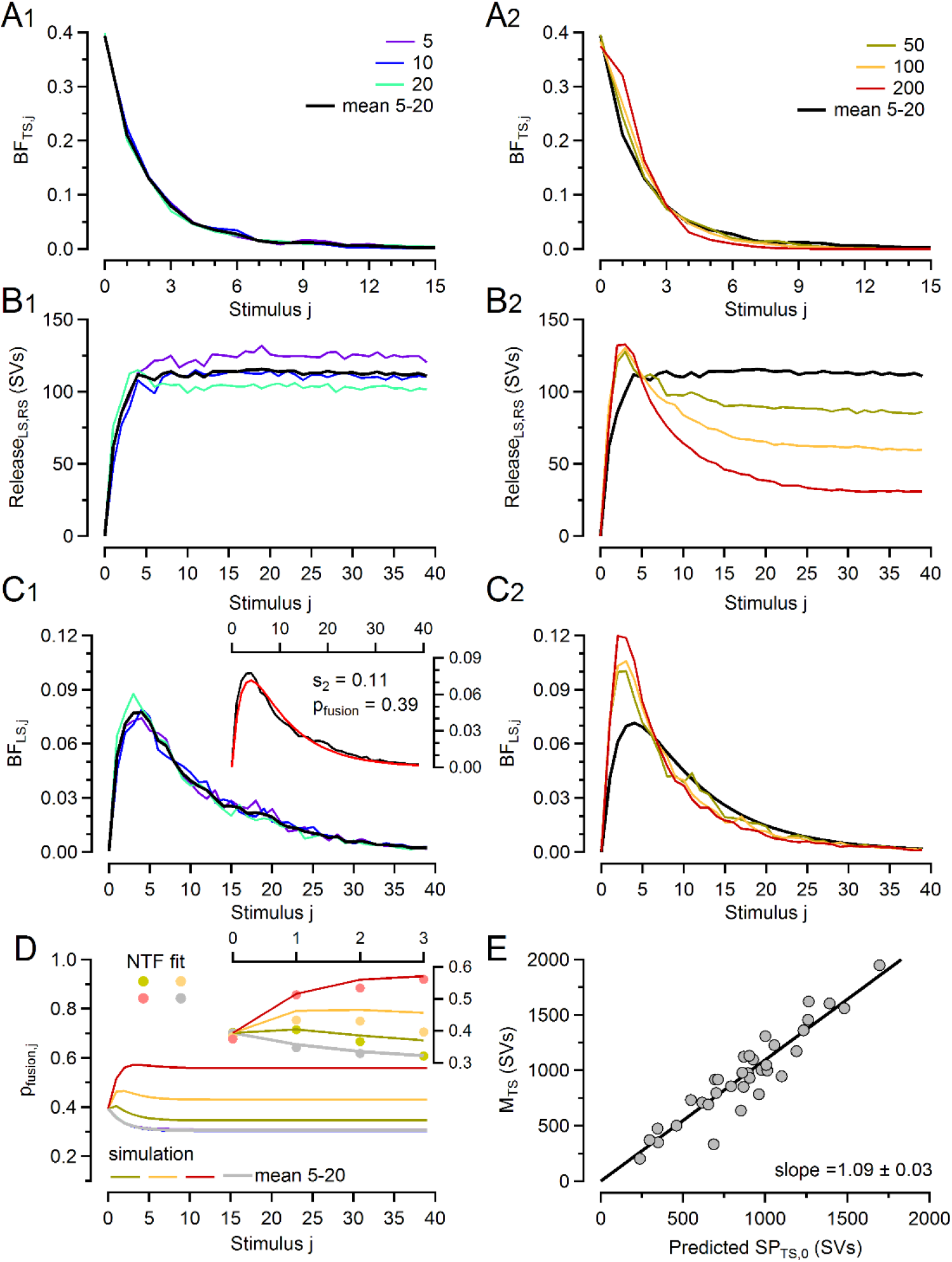
NTF decomposition analysis of 5 to 200 Hz eEPSC trains. A. Comparison of BF_TS_ reflecting the normalized release time course of pre-existing SV_TSs_ for 5, 10 and 20 Hz (**A1**) and 50, 100 and 200 Hz (**A2**) stimulation. The black traces in **A1** and **A2** represent the mean BF_TS_ for 50, 10 and 20 Hz. Individual BFs in A1 are indistinguishable from their mean.. **B.** Comparison of mean *MLS,RS* · BF_LS,RS_ reflecting the release contribution by SVs which were not tightly docked prior to stimulation, i.e. the sum of pre-existing SVLSs and newly recruited SVs, for 5, 10 and 20 Hz (**B1**) and 50, 100 and 200 Hz (**B2**) stimulation. The black traces in **B1** and **B2** represent the mean BF_LS,RS_ for 50, 10 and 20 Hz. **C.** Comparison of BFLS reflecting the normalized release time course of pre-existing SVLSs for 5, 10 and 20 Hz (**C1**) and 50, 100 and 200 Hz (**C2**) stimulation. The black traces in **C1** and **C2** represent the mean BFLS for 50, 10 and 20 Hz. The inset compares the average BFLS 5–20 Hz (black) with a fit (red) as described in Neher and Taschenberger (45) which can be used to obtain an estimate for *s*2. **D.** Simulated time course of *p_fusion_* during stimulus trains calculated according to eqn. 37. The inset compares NTF- derived *p_fusion_* estimates for the four initial eEPSCs with simulated values. **E.** Scatter plot of NTF- derived *MTS* versus *SP_TS_*_,0_ as predicted for 10 Hz eEPSC trains from *m1* and *pfusion* according to eqn. 31(see also Figure 2F). The slope of the regression line (1.09) indicates that the NTF- derived *MTS* is on average ∼9% larger.

Even though NTF analysis cannot provide initial guesses for all model parameters, it is very instrumental in constraining some of them: Two-component NTF, which is robust (45), separates release contributed by pre-existing SV_TSs_ from other contributions. BF_TS_s characteristically decay rapidly and approach zero after ∼5 APs (Fig. 3A). Their initial value represents *p_fusion_*,_1_, which amounted to 0.39 ± 0.004 when averaged over all six *f_stim_*. Provided that all pre-existing SV_TSs_ are consumed during trains, the train quantal content (*MTS*) associated with the BF_TS_ represents an estimate for *SP_TS_* at rest (*SPTS,*0), which amounted on average to 961 ± 68 SVs. The three BF_TS_s for *f_stim_* = 5–20 Hz are strikingly similar when plotted against stimulus number (Fig. 3A1). They decay exponentially indicating nearly constant *p_fusion_* throughout trains at these frequencies. BF_TS_s for ≥50 Hz deviate from this pattern (Fig. 3A2). Their second values are larger, followed by a steeper decline, which is especially prominent for 200 Hz. Time courses of *p_fusion_* during trains derived from BF_TS_ (45) indicates that *p_fusion_* slightly decreases during 5–20 Hz stimulation, while for ≥50 Hz and especially at 200 Hz, *p_fusion_* increases, consistent with the experimentally observed facilitation at high frequencies (Fig 3D).

Figure 3B shows time courses for quantal release contributed by those SVs, which were not in the TS state at stimulus onset. Again, for 5–20 Hz, these time courses are strikingly similar when plotted against stimulus number (Fig. 3B1). This is surprising, since they represent fusion of SVs which undergo at least one priming step during stimulus trains and one may expect more priming to occur during longer inter-stimulus intervals (ISIs). As pointed out previously (45) and formally proven here (eqns. 10–16), this finding is consistent with a transition of a constant fraction (*s*2) of *SP_LS_* to *SP_TS_* subsequent to an AP. At steady state, the loss from *SP_TS_* due to SV fusion is compensated by an equal number of SVs replenishing *SP_TS_*. That way, SVs are supplied to *SP_TS_* at a rate linearly increasing with *f_stim_*, which yields a frequency-invariant steady-state quantal content (*m_ss_*) at 5–20 Hz (Fig. S1B–D) (48, 49).

Three-component NTF decomposition allows the separation of release contributed by pre-existing SVLSs (Fig. 3C) from that contributed by newly recruited SVs (Fig. S2A). Provided that all pre-existing SVLSs are consumed during trains, the train quantal content (*MLS*) associated with the BFLS- represents an estimate for *SP_LS_* at rest (*SPLS,*0), which amounted on average to 1078 ± 75 SVs. BFLSs start at a value of very close zero, since pre-existing SVLSs cannot fuse during the first AP and need to undergo the LS → TS transition first. For all frequencies, BFLSs quickly increase to a maximum value after 2–3 APs before decaying exponentially to near zero. Again, the time courses of BFLSs for 5–20 Hz are nearly indistinguishable (Fig. 3C1), while those for 50–200 Hz reach higher peak values and thereafter approach zero with faster decay kinetics (Fig. 3C2). As detailed previously (45; their Fig. 8B), the average BFLS for 5–20 Hz trains can be used to obtain an estimate for *s*_2_ (0.11; Fig. 3C1 inset) which represents the fraction of *SP_LS_* transferred to *SP_TS_* subsequent to each AP. In summary, analyzing BFs provides initial estimates for key model parameters, such as initial *p_fusion_*, and the size of *SP_TS_* and *SP_LS_* at rest. Several more parameters estimates can be derived from analytical expressions regarding model predictions for e.g. *PPR* and steady-state release at low-frequency stimulation (eqns. 22–25).

### ‘Balanced priming’ accounts for frequency-invariant STD at stimulus frequencies 5–20 Hz

After initializing model parameters as described, numerical simulations reproduce steady-state depression (*Dm* = *m_ss_* / *m1*) for *f_stim_* = 0.5–20 Hz already quite well (Fig.4 D dotted trace). In the absence of stimulation, occupancies of *SP_LS_* and *SP_TS_* are determined by *k*1,0, *b*1, *k*2,0, *b*_2_ (eqns. 4, 5). During stimulation, *SP_TS_* and *SP_LS_* are partially depleted, such that quantal release decreases towards *m_ss_*, which represents a balance between SV consumption and pool replenishment (Fig. 4D). At *f_stim_* ≤2 Hz, this balance depends on the frequency-independent parameters *k*1,0, *b*1, *k*2,0, *b*2, *p_fusion_* as well as on the frequency-dependent parts of *k*_1_ and *k*2. This results in a progressive drop of *m_ss_* with increasing *f_stim_* up to ∼5 Hz (Fig. 4D). At *f_stim_* ≥5 Hz, however, release and the frequency-dependent increase of *k*_1_ and *k*_2_ above their resting values dominate. Backward (unpriming) transition rates, determined by *b*_1_ and *b*2, become negligible and the net movement of SVs along the kinetic scheme occurs nearly exclusively in forward direction (priming). The resulting steady-state occupancy of SV subpools is dominated by only four parameters: Fusion probability *p_fusion_*, total number of release sites *N_tot_* and the paameters *s*_1_ and *s*_2_ (eqn. 12), the latter representing fractions of an upstream pool converted per AP to the corresponding downstream pool. Thus, *m_ss_* tends towards a nearly frequency-independent plateau value, since both SV consumption and replenishment increase approximately linearly with frequency (‘balanced priming’) (Fig. 4D dotted trace) (48, 49). A likely mechanism for the transfer of a fixed fraction of *SP_TS_* is a linear relationship between the [Ca^2+^] signal and the priming rate constant *k*2. A further requirement is a relatively constant size of *SP_LS_* which may be achieved by a similarly balanced ES ↔ LS transition. In our modeling, we simulated this scenario by assuming for both rate constants fixed resting values (*k*_1,0_ and *k*_2,0_) and slopes 1 and 2 to describe their Ca^2+^ dependence for [Ca^2+^] > [Ca^2+^]0 (eqns. 4, 5). These parameters are related to the fractions *s*_1_ and *s*_2_ of pools transferred per AP according to eqn. 12.

**Figure 4.**
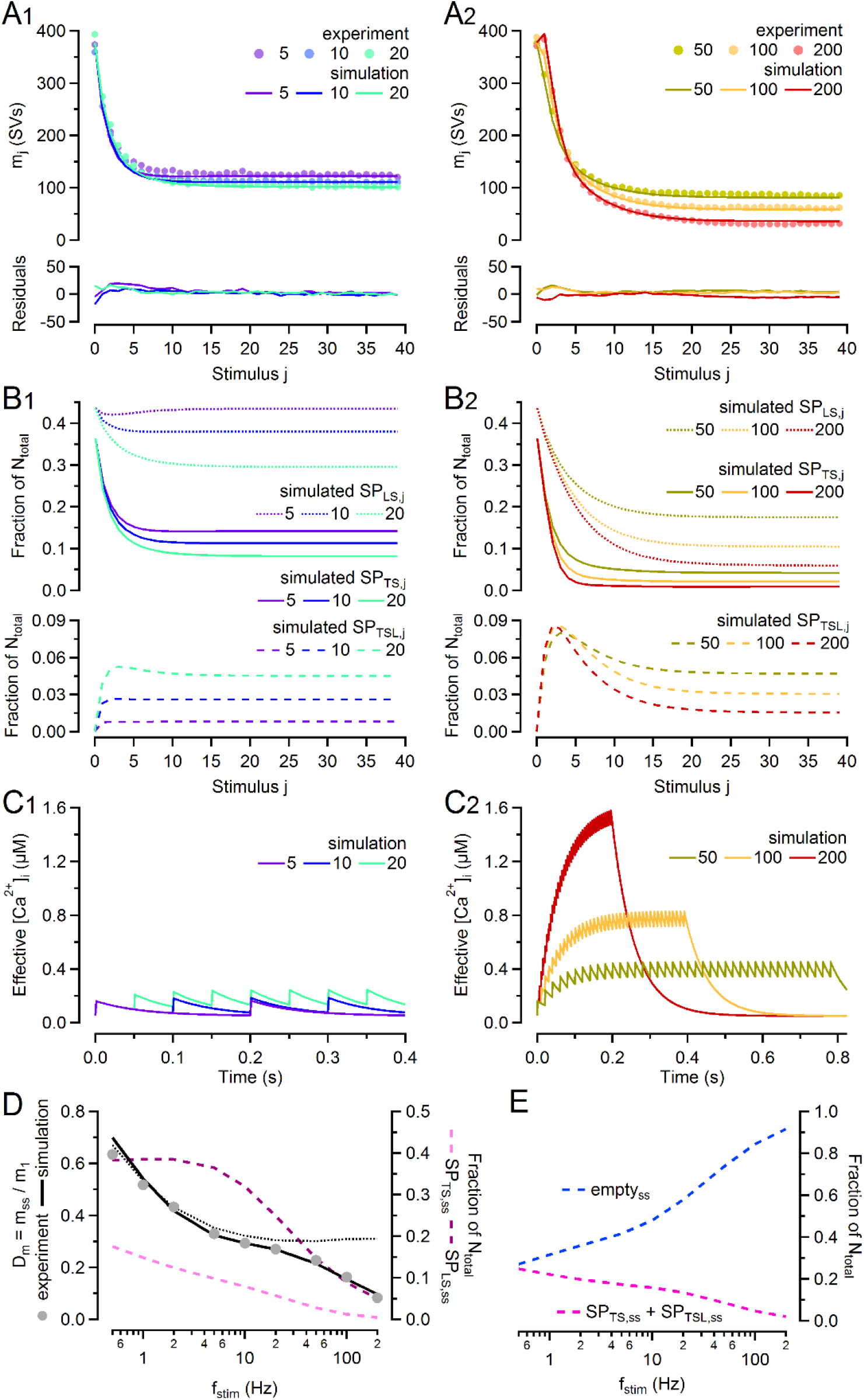
Numerical simulations of STP in response to regular 5–200 Hz stimulus trains. A. Experimental data (filled circles) and simulated *mj* values (lines) plotted against stimulus j for 5, 10 and 20 Hz (**A1**) and 50, 100 and 200 Hz (**A2**) trains. Residuals are shown in the small panels below. **B.** Time course of SV subpool occupancies immediately before AP arrival for *SP_LS_* and *SP_TS_* (top) and *SPTSL* (bottom) measured in fractions of total number of release sites (*N_tot_*) for 5, 10 and 20 Hz (**B1**) and 50, 100 and 200 Hz (**B2**) trains. Note different scaling of upper and lower panels. **C.** Time course of the effective [Ca^2+] regu^lating SV priming (eqn. 6) at high time resolution (1 ms) shown for comparison for 5, 10 and 20 Hz (**C1**) and 50, 100 and 200 Hz (**C2**) trains. Only the initial portion of the [Ca^2+^] trains are shown in **C1**. **D.** Normalized steady-state depression (*Dm* = *m_ss_* / *m*1, left axis) and steady-state occupancy of *SP_TS_* and *SP_LS_* measured in fractions of *N_tot_* (right axis) are plotted logarithmically as a function of *f_stim_*. While the steady-state occupancy of *SP_TS_* gradually declines with increasing *f_stim_*, *SP_LS_* does not substantially deplete at steady state for *f_stim_* ≤10 Hz. The dotted trace is the prediction for *Dm* by the basic model (Fig. 1B). This assumes a strictly linear *k*1(*[Ca2+]*), an empty *SPTSL* at AP arrival, no contribution of *y*(*t*) to facilitation, but a decreasing *z*(*t*) (eqn. 37), as suggested by NTF for low-frequencies. The basic model agrees quite well with the measured *Dm* (filled circles) for *f_stim_* up to 20 Hz, but clearly deviates for higher stimulation frequencies. The extended model (Fig. 1C) accurately describes data up to 200 Hz. **E.** Simulated total numbers of empty release sites (blue dashed line) and release sites occupied with a fusion-competent SV (i.e. *SP_TS_* + *SPTSL*, red dashed line) measured immediately before AP arrival at steady-state (mean of last xy values during trains) in fractions of *N_tot_* are plotted as a function of *f_stim_*.

For *f_stim_* ≥50 Hz, large deviations are evident between experimental data and this basic model, which therefore needs to be extended to reproduce STD for *f_stim_* up to 200 Hz. This also requires slight adjustments of the parameters used so far due to some overlap of high- and low-frequency features. Recovery from STD induced by conditioning ≤20 Hz stimulation is well approximated by single exponentials with time constants close to 4 s (50). This feature can be reproduced in the model by proper selection of *k*1,0 and *k*2,0.

### Multiple mechanisms of release facilitation at stimulus frequencies ≥50 Hz

Several properties of transmitter release change when stimulating at frequencies ≥50 Hz: (1) *p_fusion_* increases during trains, which is reflected in elevated second and third values of BF_TS_, followed by a more rapid decay (Figs. 3A2 and 3D). (2) Release contributed by those SVs not residing in TS prior to stimulation develops a peak around the third to fifth AP before decaying to steady-state levels lower than those at ≤20 Hz (Fig. 3B2 and 3C2). (3) Steady-state release contributed by newly recruited SVs decreases with increasing *f_stim_* causing stronger steady-state depression (Figs. 3B2 and 4D, Fig. S2A2). (4) The recovery of eEPSCs from STD develops a fast component (Fig. 5E). These changes in synapses behavior, which most likely represent interactions between release events triggered by consecutive stimuli, such as overlap of [Ca^2+] tran^sients or incomplete relaxation of internal states of the release machinery, prompted us to consider options for extending the basic sequential model (Fig. 1) in order to reproduce experimental observations.

**Figure 5.**
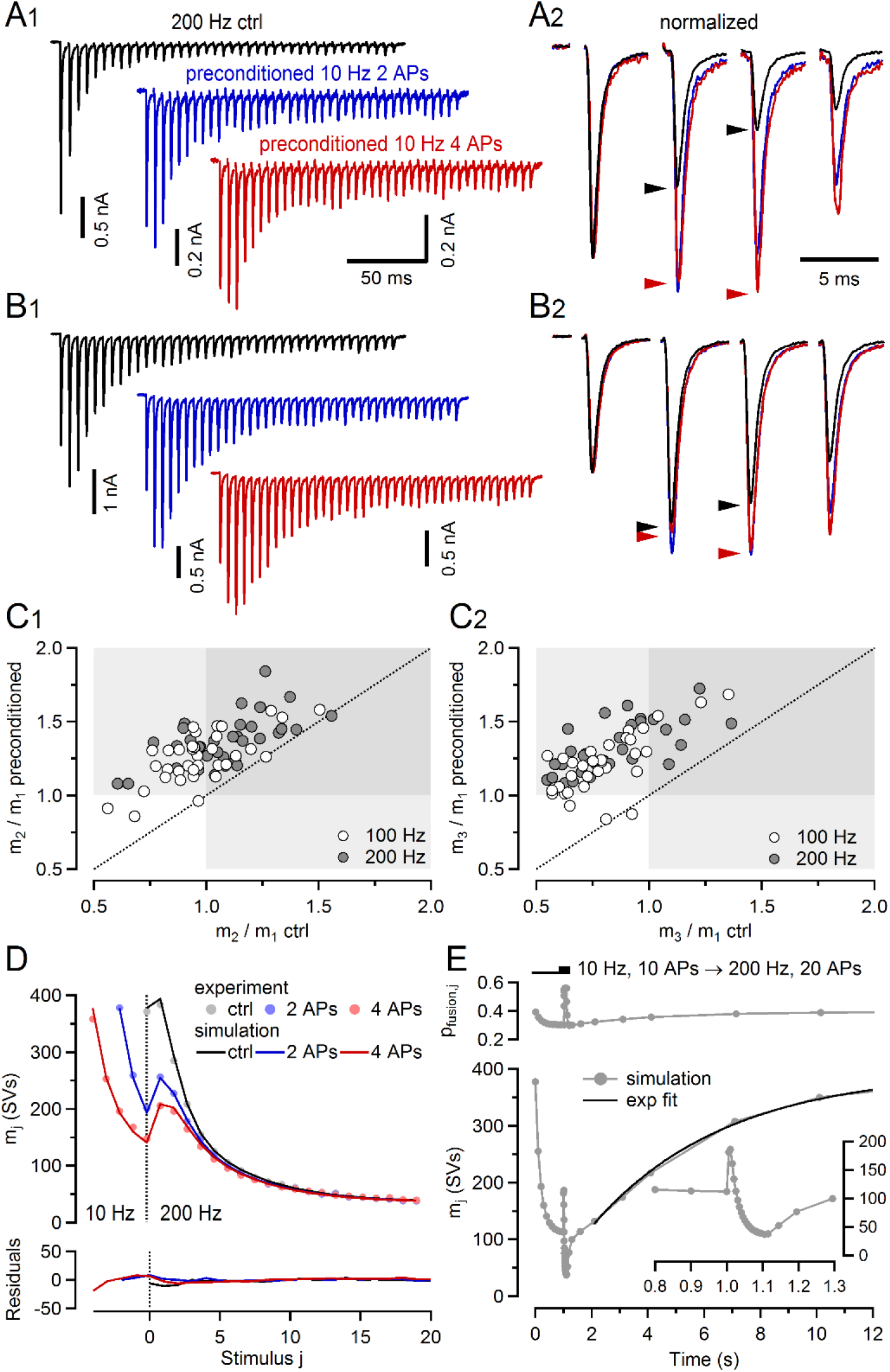
Numerical simulations of STP in response to complex stimulus train patterns. A, B. Sample 200 Hz eEPSCs recorded either without (**A1, B1,** top) or with 10 Hz pre-conditioning using 2 APs (**A1, B1,** middle) or 4 APs (**A1, B1,** bottom) in a strongly depressing (**A**) and a facilitating (**B**) synapse. 10 Hz pre-conditioning converted depression into facilitation (**A**) and augmented existing facilitation (**B**). The initial four eEPSCs are shown after normalization to the peak of eEPSC1 at an expanded time scale in **A2, B2**. Time calibration bars in **A1**, **A2** also apply to **B1**,**B2**. **C.** Ratios *m*_2_ / *m*_1_ (**C1**) and *m*_3_ / *m*_1_ (**C2**) measured after 10 Hz pre-conditioning with 4 APs plotted against the respective values obtained without pre-conditioning for all 35 synapses. Nearly all values lie above the unity line (dotted line). Values from both, 200 Hz (filled circles) and 100 Hz (open circles) eEPSC trains, are plotted. The gray shaded regions indicate ratios >1. **D.** Simulated mean *mj* values and experimental data including the conditioning responses plotted superimposed against stimulus index. Note the excellent agreement between experimental data (circles) and model predictions (solid lines). Residuals are plotted in the small panel below. The dotted vertical line marks the transition from 10 Hz conditioning to 200 Hz stimulation. **E.** Simulated mean *mj* values for a stimulus pattern consisting of 10 conditioning stimuli at 10 Hz followed by a regular 200 Hz train consisting of 20 stimuli. Thereafter, recovery of *mj* was probed with single stimulus delivered at increasing recovery intervals (0.01, 0.02, 0.05, 0.1, 0.2, 0.5, 1, 2, 3, 6, 9 s). The black line represents a single exponential fit yielding a time constant of ∼4.7 s. The inset shows the time course of *mj* around the onset of 200 Hz stimulation at higher time resolution. The simulated time course of *p_fusion_* is plotted in the small panel on top.

The increase in *p_fusion_* is likely a consequence of presynaptic *ICa* facilitation (51–53). We simulated such facilitation as predicted by NTF with an empirical model (eqns. 37–41, Fig. 3D).

Surprisingly, the increase in *p_fusion_* alone accounts only for a fraction of experimental *PPR* (see Discussion and Fig. 6A). Also, the early peak in release *MLS,RS* · BF_LS,RS_ for 200 Hz (Fig. 3B2) is not reproduced by the basic sequential model. We therefore explored additional mechanisms potentially accounting for release facilitation and very rapid priming observed at higher *f_stim_*.

**Figure 6.**
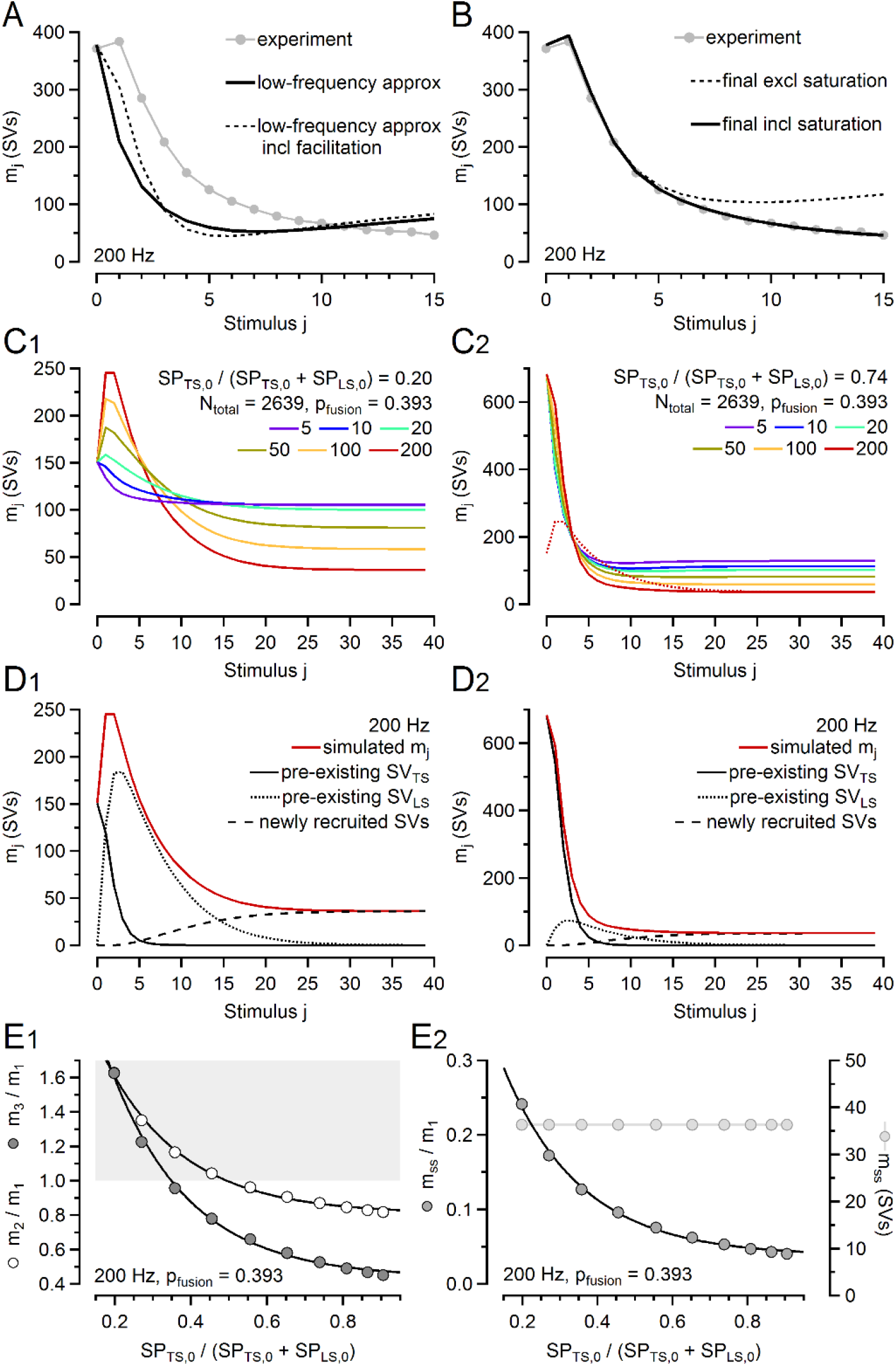
Model predictions for facilitation, depression, and STP heterogeneity A,B. Experimental data (gray symbols and lines) and several model predictions (solid and dotted black lines) for 200 Hz eEPSC trains plotted superimposed vs. stimulus index *j*. **A.** The solid trace simulates release using the basic model with constant *p_fusion_*, which is sufficient to account for experimental data up to *f_stim_* = 20 Hz. The dashed trace includes the facilitation of *p_fusion_*, as reported by NTF. Neither simulations can predict facilitation as experimentally observed for *f_stim_* = 200 Hz. **B.** Simulated 200 Hz trains using a model extended by an additional release component mediated by SVTSLs (dotted trace) and then further refined by assuming a saturating MM type relationship between [Ca^2+^] and *k*_1_ (solid trace), which accurately reproduces both PPF and STD. **C.** Time courses of *m*j in response to 5–200 Hz stimulus trains were simulated using standard values for all model parameters except for *b*_2_ which was either increased (**C1**, **D1**) or decreased (**C2**, **D2**) such that the fraction *SP_TS_*_,0_ / (*SP_LS_*,0 + *SP_TS_*_,0_) was reduced to ∼20% or enhanced to ∼74%, respectively. The red dotted trace in **C2** represents the *m*j time course for the simulated 200 Hz train shown in **C1**. **D.** Simulated contributions to release during 200 Hz stimulation by pre-existing SV_TSs_ (solid black) and by preexisting SVLSs (dotted black), or newly recruited SVs (dashed black). Simulated total release (*mj*, solid red) is shown for comparison. **E.** Ratios *m*_2_ / *m*_1_ (PPR) and *m*_3_ / *m*_1_ (**E1**) and relative steady-state depression (*m*_40_ / *m*1, **E2**) plotted versus the relative fraction of *SP_TS_*_,0_ for fourteen simulations similar to those shown in **C**,**D**. Either unpriming rate constant *b*_2_ or priming rate constant *k*_2_ for the LS ↔ TS transition were increased or decreased to generate relative *SP_TS_* fractions in the range from 0.2 to 0.9. Note, that *p_fusion_*,_1_ was kept at 0.39 and standard values were used for all other model parameters. The gray shaded region in **E1** indicates ratios >1.

For the range of experimentally used *f_stim_* values, individual AP-induced global [Ca^2+^] transients are expected to either decay nearly completely (≤20 Hz) or else summate appreciably (≥50 Hz) (Fig. 4C). A supralinear relationship between SV priming and [Ca^2+] coul^d generate release facilitation for *f_stim_* ≥50 Hz. However, the following characteristics argue against such mechanism: (*i*) The buildup of [Ca^2+] duri^ng high-frequency trains is slow, requiring ≥5 APs to reach steady-state, while facilitation is limited to only a few initial eEPSCs and is later followed by depression. (*ii*) STP of 200 Hz eEPSCs trains, which are preceded by a few stimuli delivered at 10 Hz (see below), is difficult to reproduced by the basic model. (*iii*) *PPR* (*m*_2_ / *m*1), when plotted against ISI, converts from facilitation into depression with a time constant of ∼21 ms (Fig S1D).

The frequency-dependent modulation of *p_fusion_*, as derived by NTF and predicted by the empirical model (eqns. 37–41) covers part of this decay, however cannot fully explain its degree and time course. Furthermore, with constant backward rate constants *b*_1_ and *b*2, which need to be small (<1 s^-1) in o^rder to reproduce the slow recovery of eEPSCs from STD induced by low-frequency stimulation (50), the model incorrectly predicts a long-lasting increase in eEPSC amplitude due to a persistent enhancement of *SP_TS_*. This indicates severe limitations of the basic sequential model, as used up to this point.

Considering recent ultrastructural data of presynaptic active zones (AZs) generated by time-resolved EM on samples prepared by cryofixation (39), we felt compelled to introduce a labile tightly-docked SV state (TSL) and designated SVs residing in this state as SVTSLs. We assume that each AP converts a certain fraction *κ* of *SP_LS_* into such labile SVTSLs which constitute *SPTSL* (eqns. 43–46). They have the same *p_fusion_* as SV_TSs_, however their backward transition LS ← TSL (*b*3) is more rapid than that of LS ← TS (*b*2). Thus, they contribute ‘extra’ release at the second and subsequent APs in a high-frequency train, enhancing *PPR*, but do not contribute to release, when the ISI is long longer than their mean decay time (1 / *b*3). A labile priming state can be implemented either in series between LS and TS or as a parallel branch to the LS → TS transition. For both options, model parameters can be found, which reproduce STP accurately, even when *p_fusion_* of SV_TSs_ and that of SVTSLs are constrained to have the same value. For brevity, we describe here only the version with the parallel branch (Fig. 1C). *SPTSL* and *SP_LS_* are incremented and decremented, respectively, following each AP by an amount of *κ* · *SP_LS_*. During ISIs, the time courses of SV subpool occupancies including *SPTSL* are described by the differential eqns. 43–46. By introducing this extension of a TSL together with facilitation of *p_fusion_*, we were able to numerically simulate release during high-frequency trains with high fidelity (Fig. 6B) and identified *κ* = 16% and *b*_3_ = 11.1 s^-1^ as best parameters (Table 1). These numbers are compatible with recent EM data showing that the number of SVs tightly docked at the AZ is transiently increased after an AP, but has relaxed back to normal, when flash-freezing is initiated 100 ms after the AP (39). Similar shifts between loosely and tightly docked SVs were reported to be associated with the induction of LTP in hippocampal synapses (41) as well as with beta-adrenergic modulation of parallel fiber LTP in the cerebellum (42).

### Saturation of the SV priming causes increased steady-state depression during high activity levels

Release induced by *f_stim_* ≥50 Hz obviously overstrains the processes of SV docking and priming. This leads to a progressive decline of release after the second stimulus towards a lower *m_ss_* (Figs. 4A2 and 6B) and results in a saturation of the relationship between release rate and *f_stim_* (Fig. S1C). We explored two options for modelling this: (*i*) a saturation of the priming rate with increasing *f_stim_* and (*ii*) a delayed availability of recently used sites for SV docking by introduction of a refractory release site state.

Saturation of the priming rate may occur at only the first or at both priming steps. As the simplest case, we describe here the replacement of the linear relationship between *k*_1_ and [Ca^2+^] (eqn. 4) by a Michaelis-Menten (MM) type saturating relationship (eqn. 46). This introduces a new parameter *K*0.5 which is the [Ca^2+^] at half maximum *k*1. A *K*0.5 of 280 nM and slight adjustment of other parameters adequate reproduces experimental data for all frequencies (Fig. 4A, Table 1). Note, that the simulated mean steady-state depression *Dm* = *m_ss_* / *m*_1_ exactly superimposes on experimental data over two orders of magnitude (Fig. 4D).

Having established an adequate fit to the experimental data, we then examined time courses of model quantities such as effective [Ca^2+^], *p_fusion_*, and SV subpool occupancies (Figs. 4B,C and 3D). At 10 Hz stimulation, subpool occupancies remain relatively constant during the train, except for *SP_TS_*, which rapidly depletes to a level of 0.310 times its resting value. This decay is a major cause of observed STD at 10 Hz (*m_ss_* / *m*_1_ = 0.312 ± 0.013). Relative constancy of *SP_LS_* is a consequence of balanced SV fusion and SV recruitment. At 200 Hz stimulation, both *SP_LS_* and *SP_TS_* partially deplete, and the depletion of the former is a consequence of the MM type saturation of *k*1. Depletion of SV subpools is partially compensated by an increase in *p_fusion_* (Fig. 3D). Model predictions for steady-state subpool sizes and depression are plotted against frequency in Fig. 4D,E. Simulated BFs are shown in Fig. S3. The contributions of simulated NTF components to total release at 200 Hz are shown in Fig. 6D for two cases of differing initial subpool sizes.

As an alternative to a saturation of *k*1, we introduced an empty but refractory release site state (ERS) (54–56). In this case, release sites vacated following SV fusion are not instantaneously converted into empty sites that are available for SV docking, but shifted into ERS, from which they transition with a rate constant *b*_4_ into empty sites available for SV docking (Fig. 1C2). A *b*_4_ of 0.15 s^−1^ and slight optimization of other parameters (Table 1) resulted in model predictions, which were hardly discernable from those of the alternative approach of using a MM-type saturation of *k*_1_ (Fig. 1C1).

### Accelerated eEPSC recovery after depleting high-frequency trains and STP induced by conditioning stimulation

Following high-frequency stimulation, the time course of recovery from STD exhibits an additional rapid component (13, 57). Even though recovery data were not analyzed here, the adequacy of the model can be tested against general features of recovery. As described above, the slow recovery from STD after ≤20 Hz trains (50) can be reproduced by proper selection of *k*1,0 and *k*2,0. Following stimulus trains ≥50 Hz, three processes overlap: A fast drop of *p_fusion_* from a facilitated value back to *p_fusion_*,_1_, a rapid decrease of *SPTSL*, and an increase in *SP_TS_* due to accelerated priming while [Ca^2+] is s^till elevated. The net result may either be an accelerated recovery or a transient decrease of synaptic strength, depending on the relative magnitudes of these processes and their kinetics. Ca^2+-depen^dent priming is sensitive to the incremental [Ca^2+] buil^dup during trains governed by the decay of individual AP-induced [Ca^2+] tran^sients. At ≤20 Hz, the Ca^2+- depe^ndence of *k*_1_ and *k*_2_ is relevant only in terms of the integral over [Ca^2+^] transients elicited by individual APs, i.e. large [Ca^2+] tran^sients with short duration are as effective in promoting priming as are small transients with correspondingly longer duration. At ≥50 Hz, however, [Ca^2+] tran^sients summate. During stimulus trains, the increased rate of SV priming is balanced by release. Once stimulation stops, refilling of partially depleted SV subpools continues at an increased rate until [Ca^2+^] has decayed back to [Ca^2+^]0. Lengthening the decay of the AP-induced [Ca^2+] tran^sient at the expense of its amplitude shifts a larger proportion of the priming effect into the early recovery time course immediately following the depleting stimulus train. Therefore, an amount of accelerated eEPSC recovery matching experimental data can readily be achieved in simulations by selecting an appropriate value for the model parameter *τCa* (Table 1). It should be noted, though, that experimentally observed decay time courses of global [Ca^2+] are^ often biphasic (57–59) with a rapid component much faster than the *τCa* value of 60 ms, which was appropriate for our fitting. As stated, this is of no concern when simulating low-frequency trains, for which only the time integral over the [Ca^2+] tran^sient is of relevance. However, for high frequencies, a more detailed analysis of the signal relevant for controlling Ca^2+-depen^dent priming (‘effective [Ca^2+]’), w^hich needs to consider additional aspects such as Ca^2+ buffer^s, diffusion, and morphology, may improve accuracy of model predictions.

Pronounced release facilitation can be observed at the onset of 200 Hz trains when they are preceded by a few stimuli at 10 or 20 Hz (60). The conditioning low-frequency stimulation depletes *SP_TS_*, thereby exposing the strong contribution to facilitation by the accelerated LS → TS conversion at the onset of the high-frequency train (Fig. 3B2,C2) in addition to the increase in *p_fusion_* (Fig. 3D). We analyzed 100 and 200 Hz eEPSC trains preconditioned by either 2 or 4 APs delivered at 10 Hz. Two examples for eEPSCs elicited by such stimulus pattern, which are part of our standard NTF protocol, are shown in Fig. 5A,B. Nearly all synapses showed more or less pronounced facilitation following 10 Hz pre-conditioning (Fig. 5C). The agreement between model predictions and average data is remarkable (Fig. 5D), considering that model parameters were largely determined on the basis of regular trains without preconditioning. For achieving this level of agreement, we had to increase *SP_LS_*,0, the number of SVLSs at rest, by ∼7% beyond the value predicted by NTF. All simulations shown here were performed with this slightly larger *SP_LS_*,0.

Finally, we simulated a more complex stimulus pattern consisting of 10 stimuli at 10 Hz, followed by 20 stimuli at 200 Hz and finally single stimuli delivered at various intervals to probe the time course of recovery from STD (Fig. 5E). For such stimulus pattern, the model predicts substantial depression during the 10 Hz episode (*m*_10_ / *m*_1_ = 0.301). Upon onset of 200 Hz stimulation, strong facilitation is observed (*PPR m*12 / *m*_11_ = 1.61), which quickly turns into more severe depression (*m*_30_ / *m*_1_ = 0.104). A *PPR* of 1.61 is considerably larger than the respective average *PPR* observed without preconditioning (*m*_2_ / *m*_1_ = 1.07 ± 0.03). The predicted *p_fusion_* (Fig. 5E upper panel) decreases during 10 Hz stimulation by 23%, which is followed during 200 Hz stimulation by a transient increase above *p_fusion_*,_1_ (42%). Decrease and increase in *p_fusion_* are generated by the time courses of *z* (eqns. 39, 40) and *y* (eqns. 38, 40), respectively. At the end of the 200 Hz episode, both parameters revert to their resting values as determined by the rate constants *ky* and *kz* (eqns. 40, 41; Table 1). It should be noted, that the predicted changes in *p_fusion_* are largely based on an empirical model, which reproduce NTF results (Fig 3D).

## DISCUSSION

Our analysis of STP at rat calyx of Held synapses is based on a comprehensive data set obtained under postsynaptic voltage-clamp from 35 synapses in which eEPSC trains were recorded following afferent fiber stimulation at 0.5–200 Hz. Our approach neglects delayed asynchronous release, which builds up during ISIs and slowly decays after stimulation (61). This release component is small and further developmentally reduced during postnatal synapse maturation (62). We focused on presynaptic mechanisms regulating STP by (*i*) choosing P14–16 calyx synapses in which eEPSCs are only little affected by AMPAR saturation and desensitization (63, 64), and (*ii*) by recording eEPSCs in the presence of 1 mM kyn to alleviate remaining AMPAR saturation and desensitization (63, 65). Furthermore, presynaptic feedback inhibition of release via mGluRs is reportedly low at this maturation stage (66, 67), which leaves the time course of *p_fusion_* during repetitive stimulation and the dynamic regulation of SV priming as major determinants of STP.

By analyzing this data set using a combination of non-negative tensor factorization (NTF) and conventional state modelling, we reached a number of conclusions which provide a novel view on some of the mechanisms of neurotransmitter release and STP: (1) About 80% of available presynaptic release sites are occupied at rest by primed SVs which can be either in a loosely (SVLS) or a tightly (SVTS) docked state. (2) Different initial strength and diverse STP among synapses is primarily due to variable relative abundancy of SV_TSs_ over SVLSs, while *p_fusion_* is relatively uniform across synapses. (3) Fusion-competent docked and primed SV_TSs_ have a high *p_fusion_* of ∼0.4. (4) For frequencies 5–20 Hz, ‘balanced priming’ scales steady-state release rates roughly linearly with presynaptic firing rates, thus ensuring largely frequency-invariant synaptic strength. At frequencies ≥50 Hz, additional kinetic features become apparent such as an increase in *p_fusion_* during trains, a speed-up of the priming process, and a decline of SV subpool occupancies reducing steady-state release. Before discussing these aspects individually, we summarize the assumptions underlying our analysis.

### Assumptions underlying our kinetic scheme of SV priming and fusion

For the formulation of the kinetic SV priming and fusion scheme (Fig. 1) we consider a simple sequence of steps in the buildup of a mature fusion-competent release machinery, which implies the following five main features: (1) Docking and priming of SVs occurs at a single type and a fixed number of release sites. (2) SV docking and priming steps are reversible and in dynamic equilibrium with each other at rest. (3) Release sites can be either empty or occupied by an SV residing in one of two states of functional maturation of its release machinery (referred to as SVLS and SVTS). (4) Only SVs equipped with a mature release machinery (SV_TSs_) are fusion-competent. (5) The priming rate constants *k*_1_ and *k*_2_ are Ca^2+^-dependent, while the unpriming rate constants *b*_1_ and *b*_2_ have fixed values.

### Assumptions underlying NTF decomposition of quantal release during eEPSC trains

NTF-based decomposition of eEPSC trains into release components rests on the following three assumptions: (1) Contributions to release are non-negative. (2) Release can be decomposed into distinct components, each of which represents the contribution by SVs that had been in a certain state prior to stimulation (LS, TS or undocked). (3) Forward transition rates of SV priming and fusion strongly dominate over backward transition rates in the *f_stim_* range used for NTF analysis (5–200 Hz). (4) Heterogeneity among synapses with respect to initial strength and STP characteristics is primarily due to differences in the relative abundance of SVLSs and SV_TSs_ at rest, i.e. individual synapses are endowed with variable fractions of docked SVs equipped with a mature release machinery.

NTF decomposes release during eEPSC trains into components constrained only by non-negativity. Therefore, a given NTF fit result should not be regarded as a unique solution, but rather as one out of many options consistent with both the mean eEPSC trains of all synapses examined as well as with their variability. Provided the above-mentioned assumptions hold, NTF analysis delivers good guesses for the initialization of model parameters, which together with trial-and-error parameter optimization leads to a number of important conclusions discussed below.

### A simple equation for estimating pfusion from low frequency stimulation-induced STD

When analyzing 5–20 Hz eEPSC trains, we postulated (*i*) a linear relationship between [Ca^2+] and^ priming rate constants *k*_1_ and *k*2, and (*ii*) assumed that the integral over the AP-induced [Ca^2+^] transient is constant over that *f_stim_* range. This results in ‘balanced priming’ with the average SV recruitment rate being proportional to *f_stim_*. The same conclusions can be reached, however, also with much less specific assumptions. Any process, by which an AP causes the release of a certain fraction of fusion-competent SVs and also the transition of a certain fraction of immature SVs to the fusion-competent state, will reach a frequency-independent steady state. This holds even if recruitment were non-linearly dependent on [Ca^2+] as l^ong as AP-induced [Ca^2+] tran^sients summate only little, such that each AP exerts its Ca^2+-depen^dent effect independently. This is largely the case for *f_stim_* ≤20 Hz. The choice of the exact time course of [Ca^2+^] transients has little influence on the outcome of simulations in this *f_stim_* range, because different [Ca^2+^] transients will similarly influence release as long as they have the same time integral and decay within the ISI back to [Ca^2+^]0 (eqns. 12–14).

The simplicity of conditions in the 5–20 Hz range (almost constant *p_fusion_*, balanced loss and gain for *SP_LS_*, fixed fraction of source pools transferred to downstream pools per AP), allowed us to derive a simple equation for calculating *p_fusion_*,_1_ from the two easily measurable quantities *PPR* = *m2* / *m1* and *Dm* = *m_ss_* / *m*1. In this expression (*p_fusion_*,_1_ = (1 – *PPR*)/ (1 – *Dm*); eqn. 31), the numerator recapitulates the standard use of *PPR* as an indicator for *p_fusion_* (17, 68). The equation shows, that 1 – *PPR* is actually a good estimate for *p_fusion_* in case of complete steady-state depression (*Dm* = 0), provided that all simplification made in the derivation of eqn. 31 can be applied. However, SV pool depletion is usually far from complete for 5–20 Hz stimulation for which eqn. 31 holds. Thus, the correction by the denominator is substantial. The *p_fusion_*,_1_ value obtained that way (0.43) is quite close to that derived from NTF analysis (0.396), the latter one being used in all numerical simulations. Estimates for *p_fusion_*,_1_ obtained according to eqn. 31 for individual synapses show less variability than the corresponding *m*_1_ values, which argues against *p_fusion_*,_1_ being the main source of heterogeneity in synaptic strength among synapses (Fig. 2D–F).

### Comparison to pfusion estimates derived by ‘traditional’ quantal analysis methods

For a parallel SV priming and fusion scheme, the mean *p_fusion_* is simply the weighted average of the *p_fusion_* values pertaining to individual subpools of fusion-competent SVs. The mean *p_fusion_* may be low, if ‘reluctantly releasing’ SVs (12, 21, 69, 70) contribute to release. In the context of the sequential priming scheme discussed here, NTF decomposition and modelling analysis suggest a relatively high *p_fusion_* for SV_TSs_ constituting *SP_TS_*, which represents the only fusion-competent SV subpool (Fig. 1B1). In contrast, ‘traditional’ estimates often calculate *p_fusion_* as the ratio of the quantal content of a single synaptic response over the size of the pool of readily-releasable SVs as determined by a high-frequency pool-depleting stimulus train (71). For the sequential priming scheme described here, such stimulus trains deplete not only *SP_TS_* but rather the sum *SP_TS_* + *SP_LS_* due to the rapid LS → TS transition during high-frequency stimulation. Release probability as measured by ‘traditional’ methods, therefore is the product *p_fusion_* · *SP_TS_* / (*SP_LS_* + *SP_TS_*), i.e. *p_fusion_* times the probability of a docked SV being in the tightly-docked state equipped with a mature release machinery (45). Considering a two-step SV priming scheme with only one fusion-competent state, *p_fusion_* as determined by NTF or model fitting is a quantity strictly reflecting the SV fusion process, whereas *p_fusion_* as determined by ‘traditional’ methods is a quantity depending on the fusion process as well as on the priming equilibria at rest (72, 73) (Fig. S4). This insight, if applicable to a given type of synapse, has strong implications for the interpretation of perturbations of synaptic transmission and STP by mutagenesis or pharmacological tools: An observed change in *p_fusion_* is generally interpreted as a change in the fusion machinery, involving SNARE-proteins, synaptotagmins and complexins, or else reflecting changes in the microdomain [Ca^2+] sign^al, which depends on Ca^2+ curren^ts, Ca^2+ buffer^s and coupling distances between voltage-gated Ca^2+ channe^ls (VGCCs) and Ca^2+ sensor^s for SV fusion. According to our interpretation, an observed change in a ‘traditional’ *p_fusion_* estimate may well reflect changes in the LS ↔ TS equilibrium at rest, possibly involving Munc13, DOC2, CAPS and Synaptotagmin7 (29, 32). A modulatory influence of second messengers ([Ca^2+^], DAG, PIP2) may show up in ‘traditional’ quantal analysis as a change in *p_fusion_* while NTF analysis reports it as a shift in the state of priming at rest. More precisely, the ratio of ‘traditional’ *p_fusion_* estimates over NTF-derived *p_fusion_* estimates equals *SP_TS_* / (*SP_LS_* + *SP_TS_*). Provided conditions can be found for a give type of synapse for which eqn. 31 apply, *p_fusion_*,_1_ can be estimated from *PPR* and *Dm* measured at a suitable frequency, and *SP_TS_*_,0_ is readily calculated as the ratio *m*_1_ / *p_fusion_*_,1_. Assuming further that pool estimates obtained by analyzing depleting high-frequency eEPSC trains represent the sum *SP_LS_*,0 + *SP_TS_*_,0_, then both *SP_TS_*_,0_ and *SP_LS_*,0 can be determined approximately from a small number of measurements without kinetic modelling or NTF analysis.

### Synaptic facilitation and saturation of SV priming shape the release time course during high-frequency trains

Compared to the *f_stim_* range of 5–20 Hz, for which NTF provided quite stringent constraints on model parameters, our findings regarding STP mechanisms at *f_stim_* ≥50 Hz can be interpreted in several ways. While 5–20 Hz eEPSC trains can be modelled as independently occurring release events largely unaffected by the ISIs length, consecutive release events during 50–200 Hz stimulation influence each other. They interact in various ways, through summation of global [Ca^2+] tran^sients, through kinetic limitations, e.g. saturation of priming rates, as well as via synergies, e.g. a supralinear dependence of *p_fusion_* on increasing local [Ca^2+^] due to VGCC facilitation. Together, these interactions shape synaptic facilitation and depression which can be prominently observed during 200 Hz stimulation during which initial and transient net facilitation of eEPSCs is followed by strong depression towards a small *m_ss_* (Fig. 4A2). To illustrate how a given interaction influences the release time course, we compare in Fig. 6A,B experimental data with model predictions. The basic model, which is sufficient for describing low frequencies- induced STP, fails to reproduce the 200 Hz data (Fig. 6A solid trace). Allowing *p_fusion_* to vary as predicted by NTF analysis and reproduced by the empirical *y*-*z* formalism (Fig. 3D) is insufficient to explain the extent of PPF (Fig. 6A dotted trace): Not only *m*_2_ is far below the experimental value, but even more so are *m*_3_ and *m*4. Adding the contribution of the labile tight SV state (TSL) to release leads to an adequate fit up to *m*_8_ (Fig. 6B dotted trace). However, it predicts subsequent release to rebound, due to enhanced priming by the gradual buildup of [Ca^2+] duri^ng 200 Hz trains. Only implementing saturation of *k*_1_ reproduces the experimentally observed release time course over the entire stimulation period (Fig. 6B solid trace). Alternatively, the experimental data can be simulated satisfactorily by introduction of an empty but refractory release site state (ERS) into the model (Fig. 1C2, Table 1).

### Comparison with parallel release models and past work on the calyx of Held synapse

Previous studies of release mechanisms at calyx synapses using either presynaptic voltage-clamp depolarizations or presynaptic Ca^2+ uncagi^ng for triggering SV fusion identified two kinetically distinct release components mediated by two SV subpools with differential coupling to presynaptic VGCCs (74, 75): a slowly releasing pool (*SRP*) and a fast releasing pool (*FRP*) (70, 76, 77). *SP_LS_* and *SP_TS_* are not congruent with *SRP* and *FRP*, but represent a subdivision of the *FRP*, while the contribution of the *SRP* to AP-induced release is only minor (78).

In a sequential priming scheme as used here, kinetic components may not be readily linked to specific state transitions. In contrast, it is intuitively easy to consider two kinetic components as independent contributions of two SV populations. Their intuitive tangibility explains why parallel kinetic schemes are particularly attractive. Previously, we therefore described some of the STP features, based on a much smaller data set, in terms of a parallel model consisting of a rapidly releasing SV subpool (called ‘superprimed’ SVs) and a slowly releasing one (called ‘normally primed SVs’) (19) in analogy to studies at cultured hippocampal synapses (79). ‘Superprimed’ SVs share many properties with SV_TSs_. ‘Normally primed’ SVs in the context of a parallel kinetic scheme represent fusion-competent SVs, albeit with a low fusogenicity (19). However, in the framework of the sequential scheme, proposed here, only SV_TSs_ are considered fusion-competent. Therefore, and in view of the morphological evidence for ‘loose’ and ‘tight’ coupling, we consciously avoided the previous terminology.

Release mediated by SV_TSs_ has many features in common with the release contributed by so-called ‘pre-primed’ SVs, postulated for glutamatergic synapses in the cortex (80). Likewise, the sequential scheme proposed for cerebellar parallel fiber-molecular layer interneuron synapses (25, 26) is related to the sequential scheme favored here: Both assume reversible priming steps in sequence, postulate certain fractions of upstream SV pools being transferred to a downstream pool upon AP arrival and separating contributions to release dependent on the state of SVs prior to stimulation.

### The identity of the [Ca^2+^] signal regulating the SV priming rate

For our modelling, we assumed a [Ca^2+] time^ course (‘effective’ [Ca^2+]; Tab^le 1) similar to that of the global volume-averaged [Ca^2+] as r^elevant for controlling Ca^2+-depen^dent priming. However, the identity of the [Ca^2+] sign^al which regulates SV priming is not unequivocally established. If Ca^2+-depen^dent priming depends on Munc-13, the local [Ca^2+] tran^sient similar to that triggering SV fusion may be most relevant, since Munc-13 is an integral part of the AZ (81–83). For frequencies 5–20 Hz, each AP shifts certain constant fractions of SVs from one pool to the next downstream pool. This may be taken as evidence for a local, short-lived action of the ‘effective’ [Ca^2+] tran^sients. On the other hand, Ca^2+-depen^dent priming may not be able to take advantage of higher [Ca^2+] with^in the microdomain, if the respective Ca^2+-senso^rs have high affinity and the priming rate saturates far below the peak [Ca^2+] with^in such microdomains. A complete description of Ca^2+-depen^dent priming would have to consider endogenous Ca^2+ buffer^s, which shape the time course of both the microdomain [Ca^2+]^ (^84,^ 85) as well as the global volume-averaged [Ca^2+]^ (^58,^ 59, 86–88). In view of these uncertainties, we chose to explore the influence of the [Ca^2+] time^ course with the simplest possible model, characterized by an AP-induced increment in [Ca^2+^] given by Δ[Ca^2+^] · *y(t)*, followed by a decay towards [Ca^2+^]0 with a time constant *τCa* (Fig. 4C). Both Δ[Ca^2+^] and *τCa* were free parameters during trial-and-error optimization (Table 1). Using these simplifications, we found that the exact time course of [Ca^2+] tran^sients hardly influences model predictions, as long as consecutive transients do not overlap and their integral (Δ[Ca^2+^] · *y(t)* · *τCa*) is kept constant. Our model reproduces the biphasic eEPSC recovery time course after high-frequency stimulation (13). The magnitude of the fast recovery component is larger for slower *τCa*. Thus, proper selection of *τCa* adjusts the amplitude of fast recovery to a value of 20–30% as experimentally observed (57).

Remarkably, the degree of STD at calyx synapses is relatively frequency-invariant for *f_stim_* = 5–20 Hz (48, 49). Even though this feature was recognized early (89), it is easily overlooked if STD is not characterized over the relevant frequency range, and it may be occluded by postsynaptic AMPAR saturation and desensitization. ‘Balanced priming’, which prevents *SP_LS_* from being depleted or increased at 5–20 Hz may represent a special adaptation of the SV priming machinery in calyx of Held synapses driven by a requirement for reliable transmission in the presence of a variable rate of spontaneous AP firing.

### Sources and consequences of functional diversity among individual synapses

Figure 6C, D illustrates results of two simulations, during which all model parameters were identical except for *b2*, which was either increased or decreased to obtain a relative size of *SP_TS_*_,0_ of 20% (Fig. 6C1, D1) or 80 % (Fig. 6C2, D2), respectively. We then systematically varied *b*_2_ or *k*2,0 (Fig. 6E) and plotted for simulated 200 Hz eEPSC trains the ratios *m*_2_ / *m*_1_ and *m*_3_ / *m*_1_ as a function of the relative size of *SP_TS_*_,0_ (Fig. 6E1). We observed negative correlations, reminiscent of the relationship between *PPR* and the measured *m*_1_ (Fig. 2D). A similar negative correlation was found between the simulated steady-state depression *m_ss_* / *m*_1_ and simulated *SP_TS_*_,0_ (Fig. 6E2), illustrating that the magnitude of depression strongly depends on the relative abundance of SV_TSs_ at rest. Figure 6E2 also plots the absolute *m_ss_* as a function of the relative size of *SP_TS_*_,0_ for simulated 200 Hz trains. As expected, *m_ss_* is independent of the initial distribution of SV subpool sizes at rest, since Ca^2+-depen^dent parameters of the priming scheme dominate at high frequencies over the small rate constants, which set the distribution of states at rest.

Recent studies emphasize variations in coupling distances between presynaptic VGCCs and docked SVs as a predominant mechanism generating differences in *p_fusion_* (for review see 90). Our implementation of NTF analysis, on the other hand, postulates differences in the maturation state of primed SVs as the main source of variability. It is well conceivable that a different NTF implementation, allowing several states or types of release sites to contribute to release, would come up with BFs for such states with different *p_fusion_*, as one might expect for different coupling distances. However, our assignment of variability to differences in the maturation state of primed SVs is self-consistent in the sense that eqn. 31, when applied to individual synapses, confirms that variable *p_fusion_* contributes little to their heterogeneity in synaptic strength (Fig. 2E). Of course, this does not preclude that spatial coupling between Ca^2+ entry^ and the Ca^2+ sensor^ for SV fusion may contribute to variability in synaptic strength.

The identification of molecular mechanisms causing functional synaptic diversity has recently attracted great attention due to the recognition of its importance for maximizing the capacity of information processing of neuronal networks (91) and of findings regarding the modulation of heterogeneity by astrocytes (92). Activity-induced changes in synaptic weight such as the redistribution of synaptic efficacy as a consequence of induction of LTP (93) have emerged as important elements in the description of the dynamics of neuronal networks (94, 95). The understanding of molecular mechanisms underlying such phenomena, however, has lagged behind. Such redistribution is a basic property of our model, controlled by the distribution of SVs between primed states at rest and not involving a change in *p_fusion_*. In Fig. 6C we compare simulated time courses of release of two synapses, one with a low ratio of *SP_TS_* / *SP_LS_* (Fig. 6C1), the other one with a high ratio of *SP_TS_* / *SP_LS_* (Fig. 6C2) at rest. In spite of the 4.5-fold difference in initial synaptic strength (m1 = 150 vs. 684 SVs), the cumulative release during the first 20 stimuli at 200 Hz is quite similar (2198 vs. 2695 SVs, Fig. S4B). This suggests changes in the LS ↔ TS equilibrium at rest and priming proteins, such as Munc13s, as the molecular basis of a redistribution of synaptic efficacy.

## Material and Methods

### Preparation

Juvenile, post-hearing onset (P14–16) Wistar rats of either sex were used. All experiments complied with the German Protection of Animals Act and with the guidelines for the welfare of experimental animals issued by the European Communities Council Directive. Acute brainstem slices were prepared similarly as previously described (19). See *SI Appendix* for details.

### Electrophysiology

Whole-cell patch-clamp recordings were made from principal neurons (PNs) of the MNTB as previously described (19). See *SI Appendix* for details.

### Decomposition of quantal release into distinct components using non-negative tensor factorization

Non-negative tensor factorization (NTF) of eEPSC trains was performed similarly as previously described (45) . See *SI Appendix* for details.

### Simulation of AP-evoked SV release and short-term plasticity

We used a the two-step SV priming and fusion scheme (Fig. 1) to numerically simulate SV fusion and short-term plasticity. Details of the model and approximations for steady-state conditions which were used to constrain model parameters are given in the *SI Appendix*.

## Acknowledgements

We thank Drs. Alain Marty and Stefan Hallermann for valuable discussions and comments on the manuscript, and I. Herfort for excellent technical assistance. This work was supported by the Deutsche Forschungsgemeinschaft (DFG, German Research Foundation) Cluster of Excellence EXC 2067 “Multiscale Bioimaging” (E.N.) and the DFG Collaborative Research Center 1286 “Quantitative Synaptology” (E.N.).

