## Supplemental Information for "A sequential two-step priming scheme reproduces diversity in synaptic strength and short-term plasticity"

**This PDF file includes:**

Supplementary text
Figures S1 to S4
SI References

### Material and Methods

#### *Animal Maintenance*

Juvenile, post-hearing onset (P14–16) Wistar rats of either sex were used. All experiments complied with the German Protection of Animals Act and with the guidelines for the welfare of experimental animals issued by the European Communities Council Directive. Animal health was controlled daily by caretakers. Animals were kept at 21 ± 1°C and 55% relative humidity with a 12 h/12 h light/dark cycle. Food and tap water were provided ad libitum, and cages were changed at least once a week.

#### *Slice Preparation*

Acute brainstem slices were prepared similarly as previously described (1). After decapitation, brains were dissected out quickly and immersed in ice-cold carbogenated (95% $O_2$, 5% $CO_2$) low-$Ca^{2+}$ low-$Na^+$ artificial CSF (aCSF) containing (in mM): 93 NMDG, 2.5 KCl, 0.5 $CaCl_2$, 25 glucose, 10 $MgCl_2$, 20 HEPES, 1.25 $NaH_2PO_4$, 30 $NaHCO_3$, 2 thiourea, 5 sodium ascorbate, 3 Na-pyruvate, pH 7.4 with HCl (2, 3). The brainstem was glued onto the stage of a VT1000S vibratome (Leica), and 200 µm-thick coronal slices containing the medial nucleus of the trapezoid body (MNTB) were cut. Slices were incubated for 40 min at 35°C in a chamber containing normal aCSF containing (in mM): 125 NaCl, 2.5 KCl, 2 $CaCl_2$, 1 $MgCl_2$, 25 glucose, 25 $NaHCO_3$, 1.25 $NaH_2PO_4$, 0.4 ascorbic acid, 3 myoinositol, and 2 Na-pyruvate (pH 7.4, bubbled with 95% $O_2$, 5% $CO_2$). Slices were kept at room temperature (21–24°C) and used for recordings for up to 4 h after recovery.

#### *Electrophysiology*

Whole-cell patch-clamp recordings were made from principal neurons (PNs) of the MNTB at room temperature using an EPC-10 amplifier controlled by PatchMaster software (HEKA Elektronik). Patch-pipettes pulled from borosilicate glass (Science Products) were coated with dental wax in order to reduce stray capacitance. They had an open-tip resistance of 2.5–3.5 MΩ when filled with a Cs-gluconate based solution containing (in mM): 100 Cs-gluconate, 30 TEA-Cl, 30 CsCl, 10 HEPES, 5 EGTA, 5 $Na_2$-phosphocreatine, 4 ATP-Mg, 0.3 GTP, pH 7.2 with CsOH. During experiments, slices were continuously perfused with normal aCSF solution containing 1 mM $MgCl_2$ and 2 mM $CaCl_2$ and supplemented with 5 µM strychnine, to block glycinergic IPSCs. Cells were visualized by using infrared light and oblique illumination (Dodt gradient contrast) through a 60× water-immersion objective (NA = 1.00, Olympus) using an upright BX51WI microscope (Olympus).

All experiments were performed at room temperature. A bipolar stimulation electrode was used to evoke presynaptic APs (stimulus intensity ≤20 V, 100 µs duration). Series resistance ($R_s$) was ≤8 MΩ and compensated ≥82%. Holding potential ($V_h$) and leak current were −70 mV and

≤300 pA, respectively. Sampling interval and low-pass filter settings were 20 µs and 5.0 kHz, respectively. To reduce eEPSC amplitudes for improved voltage-clamp and in order to attenuate postsynaptic AMPAR saturation and AMPAR desensitization, all experiments were performed in the continuous presence of 1 mM of the low-affinity GluR antagonist kynurenic acid (kyn) (4-6). The blocking ratio $eEPSC_{kyn}$ / $eEPSC_{ctrl}$ was measured for a subset of synapses and amounted to 0.127 ± 0.005. Voltage-clamp errors caused by remaining uncompensated $R_s$ were corrected off-line and fully compensated by applying a software correction procedure similar to that described in (7). All offline analysis of electrophysiological data and all numerical simulations were performed using Igor Pro (Wavemetrics). Differential equations were solved numerically using the fifth-order Runge-Kutta-Fehlberg algorithm implemented in Igor Pro. Original data are presented as mean ±SEM.

#### *Decomposition of quantal release into distinct components using non-negative tensor factorization*

Non-negative tensor factorization (NTF) of eEPSC trains was performed similarly as previously described (8): In a total of 35 P14–16 post-hearing calyx of Held synapses, eEPSC trains evoked by afferent-fiber stimulation using trains consisting of 40 stimuli delivered at stimulation frequencies ($f_{stim}$) of 0.5, 1, 2, 5, 10, 20, 50, 100 and 200 Hz were recorded. For NTF, however, only data within the frequency range 5–200 Hz were used. To convert eEPSC peaks into quantal content ($m$), we assumed an effective quantal size $q^*$ = −6.6 pA in the presence of 1 mM kyn in the bath (8, 9). At least three eEPSC train repetitions were analyzed for each stimulus frequency and the respective $m_j$ values for each stimulus $j$ were averaged. For the 35 synapses analyzed here, the NTF data set therefore consisted of 35 × 40 data matrices, one matrix for each of the six $f_{stim}$. NTF data were thus standard tensors consisting of one layer, except for the two highest $f_{stim}$: For 100 and 200 Hz stimulation, two additional tensors layers were obtained by pre-conditioning the synapses with either 2 or 4 stimuli delivered at a frequency of 10 Hz. During data acquisition, synapses were allowed to rest after each stimulus train to completely recover from activity-induced changes in synaptic transmission back to resting conditions (15 s following ≤10 Hz trains, 25 s following 20 and 50 Hz, 40 s following 100 and 200 Hz). The order of the stimulus trains was pseudo-randomized for each synapse to avoid a contamination of the data with systematic run-down or run-up trends. The total time of a recording session to gather all NTF data from a single calyx synapse was ~21 min.

NTF analysis was performed in two sequential steps: (1) In a first step, a two-component NTF was performed to obtain the two basefunctions (BFs) $BF_{TS}$ and $BF_{LS,RS}$, which represent release contributed by those SVs that were already in a tightly docked state ($SV_{TS}$s) prior to stimulation and of those that were not ($SV_{LS,RS}$s), respectively. Both BFs were initialized as previously described (8) and 200 iterations were used during which the goodness of fit was tracked. (2) In a subsequent step, a three-component NTF decomposition was performed providing three basefunctions $BF_{TS}$, $BF_{LS}$ and $BF_{RS}$ representing release of those SV that were either in a tightly ($SV_{TS}$s) or a loosely ($SV_{LS}$s) docked state prior to stimulation and those SVs which were in neither of these two states ($SV_{RS}$s), respectively. During this second step of analysis, the time course of $BF_{TS}$ was constrained to the result of the previous two-component NTF decomposition, and only 100 iterations were used.

The three-component NTF analysis was performed similarly as described earlier (8) with the following two modifications: (1) The estimate for the mean $M_{LS}$ was biased during iterations by shifting it at each cycle by 35% towards a pre-determined target value. This is necessary, because three-component NTF analysis does not ensure a unique separation between $M_{LS}$ and $M_{RS}$ (8). The target value was calculated the following way: Assuming that the *FRP,* as calculated by standard methods (Fig. 2C), constitutes the sum of the two components $M_{TS}$ + $M_{LS}$, we subtracted the mean $M_{TS}$ from the *FRP* to obtain an estimate for $M_{LS}$, which – provided all pre-existing $SV_{LS}$s are consumed during the trains – equals $SP_{LS,0}$. However, this estimate needs further correction, since the *FRP* is not completely depleted even at 200 Hz stimulation. Therefore, we determined this estimate, *FRP′,* for 3 frequencies (50, 100 and 200 Hz) and plotted 1 / *FRP′* vs. inter-stimulus interval (ISI). By extrapolating a regression line of this plot to ISI = 0 ms, we effectively obtain an *FRP* estimate for infinite stimulation frequency ($f_{stim}$) and assume that this approach compensates for incomplete *FRP* depletion (inset Fig. 2C) (10, 11). (2) We introduced an additional constraint on the parameter $s_2$. Again, the requirement for such a constraint derives from the fact, that NTF decomposition is compatible with a multitude of solutions, and constraints are necessary to arrive at solutions of interest. The standard NTF algorithm (8) already includes a number of general constraints, such as the requirement that $m_1$ should be very similar for all $f_{stim}$. Eqn. 25, below, can be solved for $s_2$, which we used as a constraint, which is specific for the sequential model at $f_{stim}$ = 5–20 Hz. For implementing this constraint, we fitted after each iteration a model curve for $BF_{LS}$ to the mean $BF_{LS}$ for 5–20 Hz stimulation. This fit returns an estimate for $s_2$ (see below). Deviations between the fitted $s_2$ and its value according to eqn. 25 were corrected for by compressing or expanding the time course of $BF_{LS}$ in analogy to the $p_{fusion}$ correction described in Neher and Taschenberger (8).

#### *A simple sequential kinetic scheme reproduces experimentally observed STP for frequencies up to 20 Hz*

To model the time course of synchronous AP-triggered transmitter release and the temporal pattern of STP at calyx synapses in the frequency range 1 to 20 Hz we use a kinetic scheme as shown in Fig. 1B1. We assume a fixed number of functionally identical release sites ($N_{tot}$), which at a given time *t* can either be empty ($N_e$) or occupied with a docked and primed SV. After the translocation to a release site, docking and priming of a SVs proceeds by transition through two sequential maturation states: Newly arriving SVs are initially loosely docked ($SV_{LS}$) before they mature into a tightly docked fusion competent SV ($SV_{TS}$). Thus, the entire pool of docked and primed SVs at a given time $t$ can be subdivided into a subpool of $SV_{LS}$s ($SP_{LS}$) and a subpool of $SV_{TS}$s ($SP_{TS}$), such that

$$N_{tot} = N_e(t) + SP_{LS}(t) + SP_{TS}(t) \quad (1)$$

Given the kinetic scheme illustrated in Fig. 1B, the following coupled differential equations describe temporal changes in $SP_{LS}(t)$ and $SP_{TS}(t)$ after elimination of $N_e(t)$:

$$\frac{d}{dt} SP_{LS}(t) = -(b_1 + k_1 + k_2) \cdot SP_{LS}(t) + (b_2 - k_1) \cdot SP_{TS}(t) + k_1 \cdot N_{tot} \quad (2)$$

$$\frac{d}{dt} SP_{TS}(t) = k_2 \cdot SP_{LS}(t) - b_2 \cdot SP_{TS}(t) \quad (3)$$

Here the two backward (unpriming) rate constant $b_1$ and $b_2$ have fixed values, and the two forward (priming) rate constants $k_1$ and $k_2$ are modeled as $Ca^{2+}$-dependent quantities which increase linearly with $[Ca^{2+}]$, according to:

$$k_1(t) = k_{1,0} + \sigma_1 \cdot ([Ca^{2+}](t) - [Ca^{2+}]_0) \quad (4)$$

$$k_2(t) = k_{2,0} + \sigma_2 \cdot ([Ca^{2+}](t) - [Ca^{2+}]_0). \quad (5)$$

$[Ca^{2+}]_0$ represents the resting $[Ca^{2+}]$, which was assumed to be 50 nM (12). $\sigma_1$ and $\sigma_2$ are linear slope factors characterizing the $Ca^{2+}$-dependence of the SV priming steps.

The differential equations were solved numerically during ISIs, together with a rate equation for $[Ca^{2+}](t)$, which was assumed to decay back to its resting value $[Ca^{2+}]_0$ with a rate constant $k_{Ca}$:

$$\frac{d}{dt} [Ca^{2+}](t) = -k_{Ca} \cdot ([Ca^{2+}](t) - [Ca^{2+}]_0) \quad (6)$$

For each release event, the quantal content $m_j$ of the $eEPSC_j$ triggered by stimulus $j$ was calculated as the product of $p_{rel,j} \cdot SP_{TS}(t_j)$ with both quantities evaluated immediately before stimulus arrival. Indices such as $j$, as well as $ss$ and 1 are used here and in the following to indicate stimulus index, $j$ =1 to 40, steady state and first stimulus, respectively.

To calculate $p_{fusion,j}$, we used an empirical model which requires integration of two variables $y$ and $z$, representing facilitation and a small decline in $p_{fusion,j}$ during trains, respectively, as indicated by NTF analysis (see below). Equations M5 and M6 together with M11 and M12 provide the following analytical expressions for resting values $N_{e,0}$, $SP_{LS,0}$ and $SP_{TS,0}$ at $t$ = 0:

$$N_{e,0} = N_{tot}/\left(1 + k_{1,0}/b_1 \cdot \left(1 + k_{2,0}/b_2\right)\right) \tag{7}$$

$$SP_{LS,0} = N_{e,0} \cdot k_{1,0}/b_1 \tag{8}$$

$$SP_{TS,0} = SP_{LS,0} \cdot k_{2,0}/b_2 \tag{9}$$

where $k_{1,0}$ and $k_{2,0}$ according to equations M4 and M5 are the resting values of the forward rate constants $k_1$ and $k_2$ evaluated at $[Ca^{2+}] = [Ca^{2+}]_0$.

This simple kinetic scheme of SV priming and fusion described so far reproduces quite well the experimentally observed time courses of synchronous AP-evoked release at calyx synapses for stimulus frequencies ≤20 Hz. For $f_{stim}$ ≥50 Hz, however, extensions to this scheme had to be made in order to fit experimental data without systematic deviations. These extensions (Fig. 1C) are explained in detail in in the Results section.

***Derivation of initial guesses and constraints for model parameters from steady-state conditions***

Although the simple kinetic scheme presented above only describes STP in response to $f_{stim}$ ≤20 Hz, it is nevertheless instrumental in deriving steady-state values and initial guesses for several model parameters from approximate analytical solutions. Considering that measured release time courses and NTF-derived BFs for stimulus frequencies between 5 to 20 Hz are very similar when plotted versus stimulus number $j$ (Fig. 3A), it is convenient to formulate eqns. 2–5 in terms of the ISI as the unit of time and integrating eqns. 5 and 6 over an ISI. Integrals, such as

$$\int_t^{t+\Delta t_{ISI}} k_2 \cdot SP_{LS}(t) \cdot dt = \int_t^{t+\Delta t_{ISI}} (k_{2,0} + \sigma_2 \cdot ([Ca^{2+}](t) - [Ca^{2+}]_0)) \cdot SP_{LS}(t) \cdot dt \tag{10}$$

can be approximated by

$$(\Delta t_{ISI} \cdot k_{2,0} + s_2) \cdot \overline{SP}_{LS} \tag{11}$$

where $\Delta t_{ISI}$ is the duration of the ISI, $\overline{SP}_{LS}$ is the mean value of $SP_{LS}(t)$ over the entire ISI, and

$$s_2 = \int_t^{t+\Delta t_{ISI}} \sigma_2 \cdot ([Ca^{2+}](t) - [Ca^{2+}]_0) \cdot dt \tag{12}$$

For frequencies ≤20 Hz, AP-induced $[Ca^{2+}]$ transients decay back to $[Ca^{2+}]_0$ during the ISI. In that case, $s_2$ represents the fraction of an upstream SV subpool transferred by the AP to the respective downstream pool (e.g. from $SP_{LS}$ to $SP_{TS}$). At frequencies ≥50 Hz, when $[Ca^{2+}]$ transients extend over several ISIs, this fraction is distributed over these ISIs.

An equation analogous to M12 also holds for $s_1$, the fraction of empty sites being filled during an AP. Approximating $SP_{LS}(t)$ in eqn. 10 by $\overline{SP}_{LS}$ requires that changes in $SP_{LS}$ during one ISI are relatively small. This requirement is satisfied quite well at the steady-state of eEPSC trains and for values of $s_1$ and $s_2$ of ~0.1. Constants, such as $b_1$ and $b_2$, as well as the constant parts of $k_1$ and $k_2$, i.e. $k_{1,0}$ and $k_{2,0}$, show up in the low-frequency approximation as products of $\Delta t_{ISI}$ and their respective values (eqn. 11). Performing these integrations over an ISI, we arrive at the following difference equations for the changes in SV subpool sizes ($\Delta SP_{LS}$ and $\Delta SP_{TS}$) during one ISI:

$$\begin{aligned}\Delta SP_{LS} = & -\left[\Delta t_{ISI} \cdot \left(b_1 + k_{1,0} + k_{2,0}\right) + s_1 + s_2\right] \cdot \overline{SP}_{LS} \\ & +\left[\Delta t_{ISI} \cdot \left(b_2 - k_{1,0}\right) - s_1\right] \cdot \overline{SP}_{TS} + \left[\Delta t_{ISI} \cdot k_{1,0} + s_1\right] \cdot N_{tot}\end{aligned} \tag{13}$$

$$\Delta SP_{TS} = \left(\Delta t_{ISI} \cdot k_{2,0} + s_2\right) \cdot \overline{SP}_{LS} - (\Delta t_{ISI} \cdot b_2 + p_{rel}) \cdot \overline{SP}_{TS} \tag{14}$$

In eqn. 14 we include as a decrement the number of $SV_{TS}$s fusing in response to an AP ($m = p_{fusion} \cdot \overline{SP}_{TS}$), such that $\Delta SP_{TS}$ represents the total change in $SP_{TS}$ during one ISI. Here we introduce another simplification because the number of $SV_{TS}$s fusing ($m$) is actually proportional to the product of $p_{fusion}$ with the size of $SP_{TS}$ at the time of arrival of stimulus $j$ ($SP_{TS,j}$), i.e. immediately before the beginning of the subsequent ISI. However, again the relative magnitude of this error $(SP_{TS,j} - \overline{SP}_{TS})/\,\overline{SP}_{TS}$ is small at steady-state.

Solving for the mean steady-state size of SV subpools $SP_{LS}$ ($\overline{SP}_{LS,ss}$) and $SP_{TS}$ ($\overline{SP}_{TS,ss}$) by setting $\Delta SP_{LS}$ and $\Delta SP_{TS}$ in eqns. 13 and 14 to 0, we obtain:

$$\overline{SP}_{LS,ss} = N_{tot} \cdot (\Delta t_{ISI} \cdot k_{1,0} + s_1) \cdot (\Delta t_{ISI} \cdot b_2 + p_{fusion})/S \tag{15}$$

and

$$\overline{SP}_{TS,ss} = N_{tot} \cdot (\Delta t_{ISI} \cdot k_{2,0} + s_2) \cdot (\Delta t_{ISI} \cdot k_{1,0} + s_1)/S \tag{16}$$

where

$$S = \left(\Delta t_{ISI} \cdot b_2 + p_{fusion}\right) \cdot \left(\Delta t_{ISI} \cdot \left(b_1 + k_{1,0} + k_{2,0}\right) + s_1 + s_2\right) \tag{17}$$

$$-(\Delta t_{ISI} \cdot k_{2,0} + s_2) \cdot (\Delta t_{ISI} \cdot (b_2 - k_{1,0}) - s_1)$$

Equation M16 can used to calculate the relative steady-state depression of release during eEPSC trains, defined as

$$D_m = m_{ss}/m_1 \approx p_{fusion,ss}/p_{fusion,1} \cdot \overline{SP}_{TS,ss}/SP_{TS,1}. \tag{18}$$

It represents a good estimate for $D_m$ $in$ the frequency range 1 to 20 Hz, once the parameters involved are known. In the next paragraph we describe how most of these can be determined form the experimental data.

***Prediction of readily measurable quantities at stimulus frequencies of 5 to 20 Hz***

For $f_{stim}$ = 5–20 Hz, $D_m$ approaches an asymptotic value because the terms $\Delta t_{ISI} \cdot k_{1,0}$ and $\Delta t_{ISI} \cdot k_{2,0}$ are smaller by an order of magnitude than $s_1$ and $s_2$. In addition, $\Delta t_{ISI} \cdot b_1$ and $\Delta t_{ISI} \cdot b_2$ are much smaller than the products of $\Delta t_{ISI}$ and $k_{1,0}$ and $k_{2,0}$. Thus, SVs priming proceeds predominantly from left to right in the schemes of Fig. 1. Neglecting the small terms, one arrives at two very simple equations for the increments in SV subpool sizes in analogy to M13 and M14

$$\Delta SP_{LS} = -(s_1 + s_2) \cdot \overline{SP}_{LS} - s_1 \cdot \overline{SP}_{TS} + s_1 \cdot N_{tot} \tag{19}$$

$$\Delta SP_{TS} = s_2 \cdot \overline{SP}_{LS} - p_{fusion} \cdot \overline{SP}_{TS} \tag{20}$$

These equations, together with

$$\overline{N}_{e,ss} + \overline{SP}_{LS,ss} + \overline{SP}_{LS,ss} = N_{tot} \tag{21}$$

define the steady-state values of the SV subpool sizes ($\overline{SP}_{LS,ss}$ and $\overline{SP}_{TS,ss}$) according to:

$$\overline{SP}_{LS,ss} = N_{tot}/s_2/\left(\frac{1}{p_{fusion,ss}} + \frac{1}{s_1} + \frac{1}{s_2}\right) \tag{22}$$

$$\overline{SP}_{TS,ss} = N_{tot}/p_{fusion,ss}/\left(\frac{1}{p_{fusion,ss}} + \frac{1}{s_1} + \frac{1}{s_2}\right) \tag{23}$$

The quantal content of the steady-state eEPSC ($m_{ss}$) can be approximated by

$$m_{ss} \approx \overline{SP}_{TS,ss} \cdot p_{fusion,ss} = N_{tot} / \left( \frac{1}{p_{fusion,ss}} + \frac{1}{s_1} + \frac{1}{s_2} \right) \quad (24)$$

Equations 22–24 and other readily available data were used to derive initial guesses for model parameters.

A simpler form for calculating the mean steady-state size of $SP_{LS}$ is obtained by setting $\Delta SP_{TS}$ in eqn. 20 to 0:

$$\overline{SP}_{LS,ss} = p_{fusion} \cdot \overline{SP}_{TS,ss} / s_2 \quad (25)$$

By considering that $p_{rel} \cdot \overline{SP}_{TS,ss} = m_{ss}$ and $m_{ss} = m_1 \cdot D_m$, and introducing $D_{LS}$, the relative occupancy of $SP_{LS}$ at steady state ($D_{LS} = \overline{SP}_{LS,ss} / SP_{LS,1}$), we obtain:

$$SP_{LS,1} = \frac{m_1}{s_2} \cdot \frac{D_m}{D_{LS}} \quad (26)$$

Similarly, a simple expression for the paired-pulse ratio (*PPR*), calculated from the two initial eEPSCs in a train, can be obtained as:

$$PPR = \frac{m_2}{m_1} = \frac{p_{fusion,2} \cdot SP_{TS,2}}{m_1} \quad (27)$$

Applying eqn. 20 to eliminate $SP_{TS,2}$, we obtain:

$$PPR = p_{fusion,1} \cdot r_p \cdot (SP_{TS,1} \cdot \left(1 - p_{fusion,1}\right) + s_2 \cdot SP_{LS,1}) \cdot \frac{1}{m_1} \quad (28)$$

Here, the parameter $r_p = p_{fusion,2} / p_{fusion,1}$ was introduced, which is very close to 1 for low-frequency stimulation. Considering that $p_{fusion,1} \cdot SP_{TS,1} = m_1$, and using eqn. 26 to eliminate $SP_{LS,1}$ we obtain

$$PPR = r_p \cdot (1 - p_{fusion,1} \cdot (1 - \frac{D_m}{D_{LS}})) \quad (29)$$

This equation can be used to calculate $p_{fusion,1}$:

$$p_{fusion,1} = (1 - \frac{PPR}{r_p})/(1 - \frac{D_m}{D_{LS}}) \quad (30)$$

Because $r_p$ is very close to 1 for 10 Hz stimulation, and because only little reduction in the size of $SP_{LS}$ is observed during trains at 10 Hz (Fig. 4B1), an approximate expression for $p_{fusion}$, assuming $r_p \approx 1$ and $D_{LS} \approx 1$, is simply:

$$p_{fusion,1} \approx (1 - PPR)/(1 - D_m) \quad (31)$$

In conclusion, provided that $p_{fusion,2}$ is close to $p_{fusion,1}$ and that $SP_{LS,ss}$ is close to $SP_{LS,1}$, eqn. 31 can be used to calculate $p_{fusion,1}$ from two readily determined experimental quantities: the *PPR* and the relative steady-state depression $D_m$. It should be noted, that the stated conditions are fulfilled at the calyx synapse for 10 Hz, but not necessarily at other types of synapses and at the same frequency.

***Estimates for parameters $s_1$ and $s_2$ .***

As shown previously (8; their Fig. 8), eqns. 13 and 14 can be used to simulate the time course of $BF_{LS}$ as a function of the two model parameters $p_{fusion}$ and $s_2$, the latter being designated as α in that publication. Assuming that $p_{fusion}$ is constant and known from two-component NTF, we used this possibility to obtain an estimate for $s_2$ by fitting simulated $BF_{LS}$, to the $BF_{LS}$, as obtained from a three-component NTF. An alternative estimate can be obtained by solving eqn. 26 for $s_2$, assuming a value close to 1 for $D_{LS}$, as suggested by NTF. The two methods provide very similar values close to 0.1.

Given this estimate for $s_2$ eqn. 24 can be solved for $s_1$:

$$s_1 = m_{ss} \Big/ \left( N_{tot} - m_{ss} \cdot \left( \frac{1}{p_{fusion,ss}} + \frac{1}{s_2} \right) \right) \quad (32)$$

Except for $N_{tot}$, all quantities on the right-hand side of this equation are known. We noticed that model fits do not vary in a major way, when using $s_1$ calculated with M32 and setting $N_{tot}$ to values which result in 20% to 30% empty release sites at rest. Thus, we calculated the initial guess for $s_1$ according to eqn. 32 with $N_{tot}$ set to $1.25 \cdot (M_{LS} + M_{TS})$.

***Initial guess values for model parameters describing the resting state***

In order to define a complete set of initial guess values for model fitting, four more parameters, $k_{1,0}$, $k_{2,0}$, $b_1$ and $b_2$, describing the resting state, must be determined. Eqns. 7–9 provide two constraints on the choice of these parameters. The choice of $k_{1,0}$ strongly influences the time course of recovery from depression induced by conditioning stimulation. It was therefore used as a free parameter during model fitting, and adjusted to yield a time constant of recovery from STD following low-frequency stimulation of 4–6 s (13). In addition, we found that at low frequencies, the slope of the relationship between steady-state depression and $f_{stim}$ was fitted appropriately if the characteristic frequency for the second priming step (ratio $k_{2,0}$ / $s_2$) was chosen somewhat lower than that of the first one (ratio $k_{1,0}$ / $s_1$). With

$$k_{2,0}/s_2 = 0.5 \cdot k_{1,0}/s_1, \tag{33}$$

$SP_{LS,0} = M_{LS}$, $SP_{TS,0} = M_{TS}$ and eqn. 1, we get from eqn. 8:

$$b_1 = N_{e,0} \cdot k_{1,0}/SP_{LS,0}. \tag{34}$$

From eqn. 33:

$$k_{2,0} = k_{1,0} \cdot s_2/s_1 \tag{35}$$

From eqn. 9:

$$b_2 = SP_{LS,0} \cdot k_{2,0}/SP_{TS,0} \tag{36}$$

It is important to note that eqns. 8–36 are approximations, based on strict linearity of the system, such as constancy of certain parameters and linear dependence of priming rate constants on [$Ca^{2+}$] (but see Discussion on this requirement). Nevertheless, we found that assigning initial guess values derived this way to the model parameters resulted in good fits to experimental data obtained with 1–20 Hz stimulation even without further parameter optimization (see Results section). Yet, most of them were used as free parameters in subsequent trial and error fitting.

***Local [$Ca^{2+}$] transients and release probability***

Release per action potential is a power function of the local [$Ca^{2+}$] 'seen' by the $Ca^{2+}$ sensor for release (14, 15). We therefore determined $p_{fusion,j}$, the release probability at arrival of the $j^{th}$ AP, according to

$$p_{fusion,j} = p_{fusion,1} \cdot {y_j}^{4.5} \cdot z_j \tag{37}$$

with y≥1 and z≤1. Here, $p_{fusion,1}$ designates the fusion probability for the first eEPSC in a train (Table 1), $y_j$ accounts for changes in local [$Ca^{2+}$] during repetitive stimulation ($y_j = [Ca^{2+}]_j/[Ca^{2+}]_1$), likely due to presynaptic $Ca^{2+}$ current facilitation (16, 17), and/or saturation of local $Ca^{2+}$ buffers (18-20), and *z* accounts for a reduction of $p_{fusion}$ during repetitive stimulation, which in our simulations was relatively small. A candidate mechanism generating a small decrease in mean $p_{fusion}$ during trains could be a slightly non-uniform $p_{fusion}$ among all $SV_{TS}$s of a given synapse.

Both variables $y_j$ and $z_j$ were initialized to 1 at the onset of a stimulus train. The variable *y* was incremented after each AP by

$$y_{inc} = y_{inc,1} \cdot (y_{max} - y_j) \tag{38}$$

and *z* was decremented by

$$z_{dec} = z_{dec,1} \cdot (z_j - z_{min}) \tag{39}$$

During ISIs, both *y(t)* and *z(t)* were solved numerically, together with other model parameters (see eqns. 2, 3), according to

$$\frac{d}{dt}\, y(t) = (1 - y(t)) \cdot k_y \tag{40}$$

$$\frac{d}{dt}\, z(t) = (1 - z(t)) \cdot k_z \tag{41}$$

The parameters describing *y(t)*, $y_{inc,1}$, $y_{max}$ and $k_y$, were initialized with previously published values (21) and adjusted by trial and error fitting. The parameters describing *z(t)*, $z_{dec,1}$, $z_{min}$ and $k_z$, were initialized to values, which would produce a decrease of $p_{fusion}$ of ≤15% during the first few APs at 10 Hz. All six parameters of the *y*-*z* formalism were then adjusted during trial-and-error fitting to reproduce the time course of $p_{fusion}$ as derived from NTF analysis (Fig. 3D). The $p_{fusion}$ time courses show a small negative trend for the 5–20 Hz average trace due to a decrease in the *z* variable and quite prominent facilitation for 200 Hz due to an increase in the *y* variable. Reliable NTF-derived $p_{fusion}$ estimates can only be obtained for the initial four eEPSCs from $BF_{TS}$. Later during trains, $BF_{TS}$ is very small, such that estimates are unfortunately dominated by random fluctuations. Once all six parameters of the *y*-*z* formalism were determined on the basis of the $BF_{TS}$, they were held constant during all numerical simulations of release and STP.

***Extensions of the 'Simple' sequential kinetic scheme to reproduce high frequency features***

In order to simulate the experimentally observed time courses at $f_{stim}$ ≥50 Hz, the linear model as presented above (Fig.1B) had to be extended. This can be achieved in several ways, as explained in the Results section. In our preferred variant, described here in detail, we introduced an additional fusion-competent SV state, which is similar to state TS but labile (TSL, Fig. 1C). SVs populating this state ($SV_{TSL}$s) after an AP rapidly relax to state LS due to a relatively high backward rate constant $b_3$. The experimentally observed saturation of release at high $f_{stim}$ (suppl. Fig. 1C) can be described in two alternative ways. Here, for completeness, we describe how both extensions were handled, although most of the results presented were obtained by simply introducing Michaelis-Menten (MM) like saturation of $k_1$ according to:

$$k_1(t) = \left(k_{1,0} + \sigma_1 \cdot ([Ca^{2+}](t) - [Ca^{2+}]_0)\right) / \left(1 + ([Ca^{2+}](t) - [Ca^{2+}]_0)/K_{0.5)}\right) \quad (42)$$

Alternatively, an additional empty and refractory state (ERS) can be introduced as shown in Fig. 1C2). The extended scheme, including the ERS, is described in analogy to eqns. 1–6 by:

$$\begin{aligned}\frac{d}{dt}SP_{LS}(t) = &-(b_1 + k_1 + k_2) \cdot SP_{LS}(t) + (b_2 - k_1) \cdot SP_{TS}(t) \\ &+(b_3 - k_1) \cdot SP_{TSL}(t) - k_1 \cdot N_{er}(t) + N_{tot} \cdot k_1\end{aligned} \quad (43)$$

$$\frac{d}{dt}SP_{TS}(t) = k_2 \cdot SP_{LS}(t) - b_2 \cdot SP_{TS}(t) \quad (44)$$

$$\frac{d}{dt}SP_{TSL}(t) = -b_3 \cdot SP_{TSL}(t) \quad (45)$$

$$\frac{d}{dt}N_{ERS}(t) = -b_4 \cdot N_{ERS}(t) \quad (46)$$

The differential equations M43–M46 were solved numerically during ISIs. At the time of APs, increments were applied to SV subpools: Release was calculated as ($SP_{TS}(t)$ + $SP_{TSL}(t)$) · $p_{fusion}$ and $N_e(t)$ or else $N_{ERS}(t)$, was incremented by the same amount. $SP_{TS}(t)$ and $SP_{TSL}(t)$ were decremented by their respective contributions to release.

The two parameters describing $SP_{TSL}(t)$ were *κ,* the fraction of $SP_{LS}$ transferred to $SP_{TSL}$ per AP and a backward rate constant $b_3$. Both parameters were determined by trial-and-error fitting, aiming at a correct representation of amount and time course of quantal release. For convenience, the same program code can be used for both model options of handling release saturation. For the case of Michaelis-Menten-type saturation $b_4$ can be set to a very high value (≥5000) to prevent filling of ERS while adjusting $K_{0.5}$. Alternatively, $K_{0.5}$ can be set to 10 M while adjusting $b_4$.

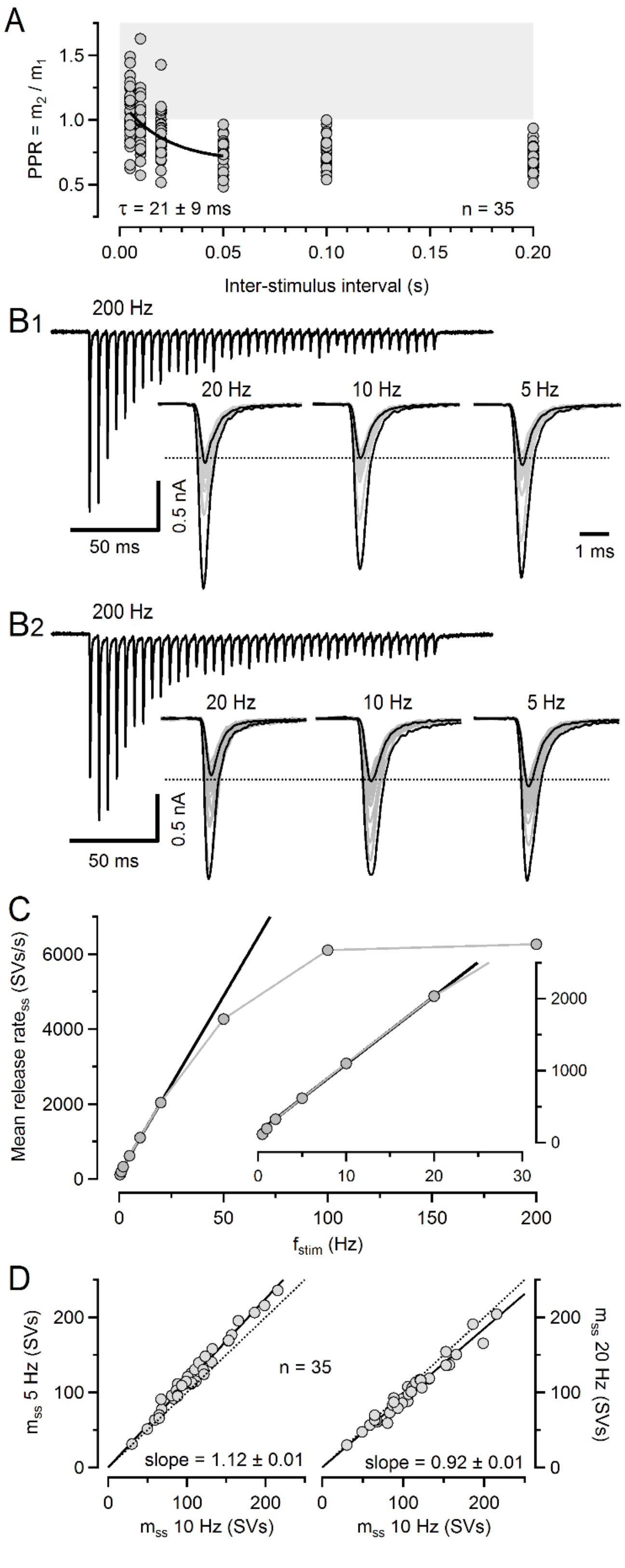
A
PPR = m2 / m1
1.5
1.0
0.5
τ = 21 ± 9 ms
n = 35
0.00
0.05
0.10
0.15
0.20
Inter-stimulus interval (s)
B1
200 Hz
20 Hz
10 Hz
5 Hz
0.5 nA
50 ms
1 ms
B2
200 Hz
20 Hz
10 Hz
5 Hz
0.5 nA
50 ms
C
Mean release rate_ss (SVs/s)
6000
4000
2000
0
2000
1000
0
0
10
20
30
0
50
100
150
200
f_stim (Hz)
D
m_ss 5 Hz (SVs)
200
100
0
n = 35
slope = 1.12 ± 0.01
slope = 0.92 ± 0.01
m_ss 20 Hz (SVs)
m_ss 10 Hz (SVs)
m_ss 10 Hz (SVs)

**Fig. S1. Characterization of *PPR*s and steady-state release.**
**A.** *PPR*s = $m_2$ / $m_1$ plotted vs. ISI for all 35 synapses. The solid line represents an exponential fit to the data in the ISI range from 5–50 ms yielding a time constant of 21 ± 9 ms.
**B.** Example eEPSC trains recorded in a depressing (**B1**) and a facilitating (**B2**) synapse stimulated at 200 Hz (top panels) and 20, 10 and 5 Hz (bottom panels). For 5–20 Hz trains, eEPSCs are shown superimposed and $eEPSC_2$ to $eEPSC_{29}$ are shown in gray. Note the similar $eEPSC_{ss}$ amplitudes for 5–20 Hz stimulation (dotted lines).
**C.** Mean steady-state release rate ($m_{40} \cdot f_{stim}$) plotted versus $f_{stim}$. The solid black line represents a linear regression to the $f_{stim}$ range 2–20 Hz, which is shown at an expanded abscissa in the inset.
**D.** Scatter graphs of $m_{ss}$ for 5 Hz (left) and for 20 Hz (right) versus the corresponding $m_{ss}$ for 10 Hz stimulation for all 35 synapses. Note the close proximity of linear regressions to the identity lines. On average, $m_{ss}$ was only 12% larger or only 8% smaller for 5 and 20 Hz, respectively, when compared to the corresponding 10 Hz $m_{ss}$ values.

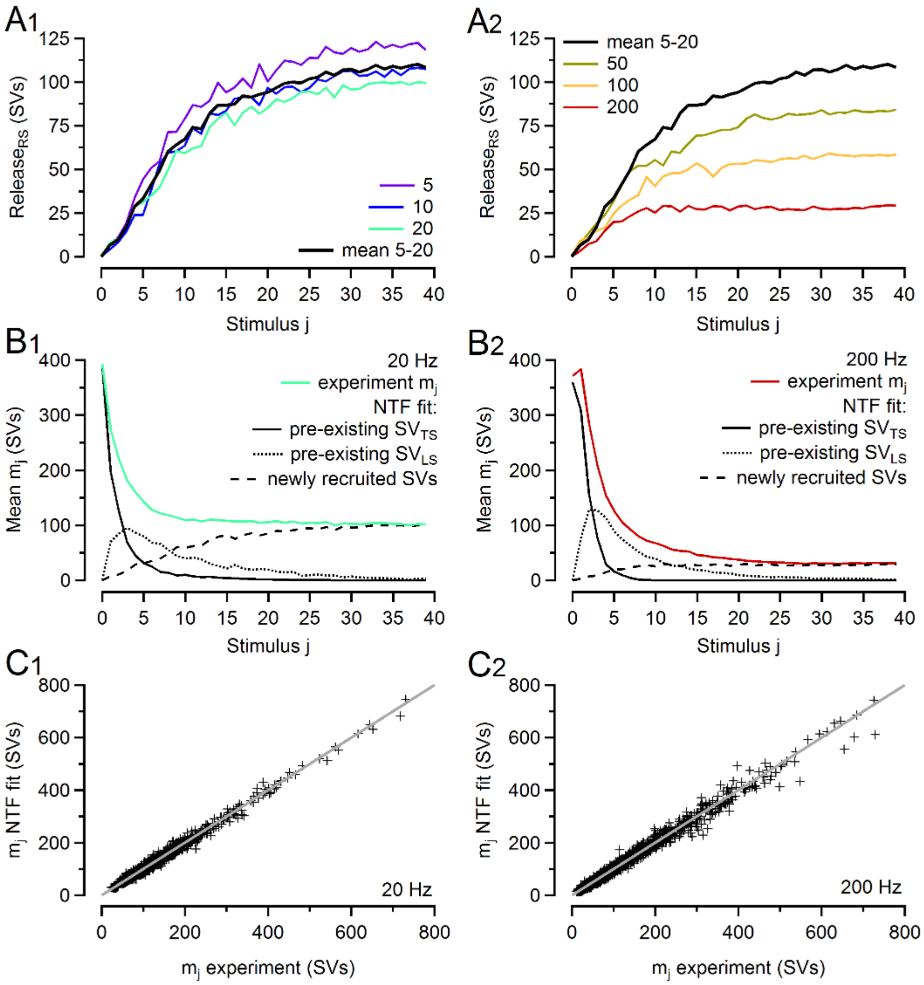


**Fig. S2. Three-component NTF decomposition analysis.**

**A.** Contributions to release by newly recruited SVs (mean $M_{RS} \cdot BF_{RS,j}$) for low (**A1**) and high (**A2**) $f_{stim}$. The black traces in **A1** and **A2** represent the mean time course for 5–20 Hz. The time courses represent release contributed by SVs, which dock to empty sites during trains and undergo the sequence of two-step priming and fusion. They start in a sigmoid fashion and approach a steady-state.

**B.** Mean $m_j$ values (solid colored traces) versus stimulus index j for 20 Hz (**B1**) and 200 Hz (**B2**). NTF fit derived mean release components contributed by either pre-existing $SV_{TS}$s (mean $M_{TS} \cdot BF_{TS,j}$, solid black line), pre-existing $SV_{LS}$s (mean $M_{LS} \cdot BF_{LS,j}$, dotted black line) and newly recruited SVs (mean $M_{RS} \cdot BF_{RS,j}$, dashed black line) are shown superimposed. Note that all pre-existing $SV_{TS}$s and all pre-existing $SV_{LS}$s are nearly completely consumed within the initial 8 and 30 APs, respectively. For stimuli ≥30, release is nearly exclusively supported by newly-recruited SVs.

**C.** Scatter plot of individual $m_j$ for all stimuli $j$ in all synapses as estimated by NTF decomposition analysis versus the respective experimentally measured values for 20 Hz (**C1**) and 200 Hz (**C2**). Note that the values are very close to the identity line and approximately symmetrically distributed around it (light gray).

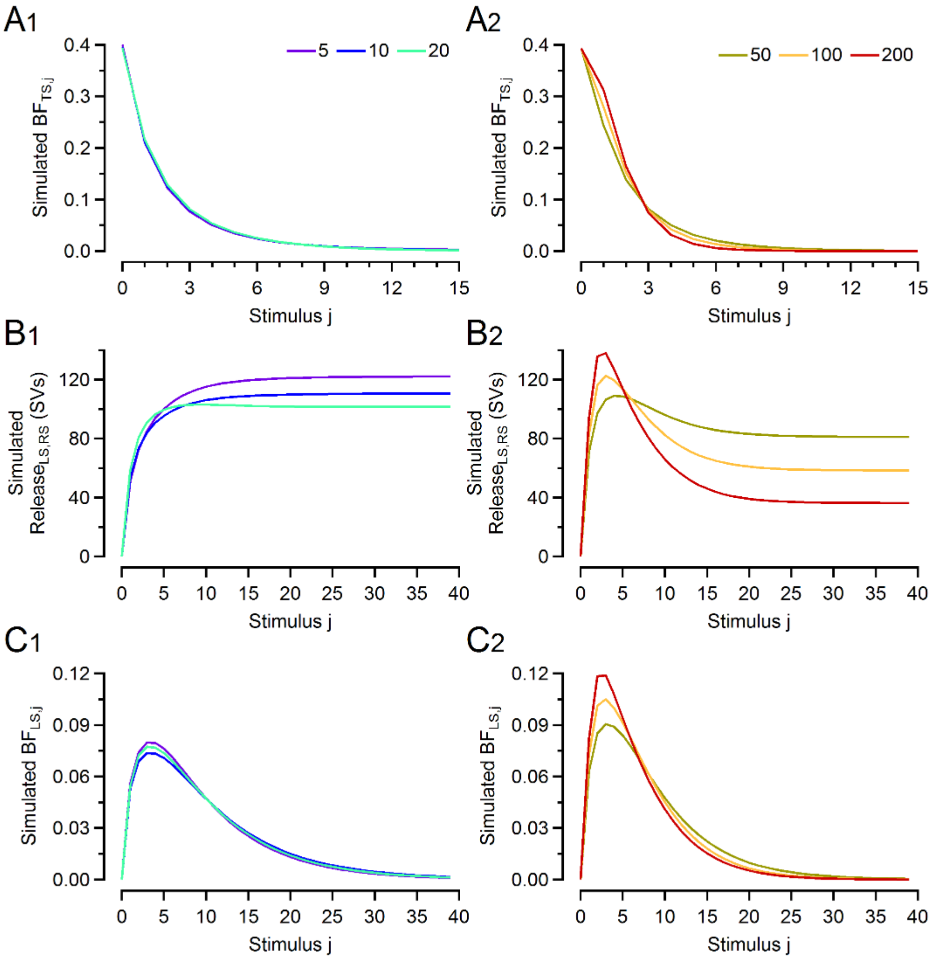


**Fig. S3. Model-derived predictions for NTF basefunctions.**
**A.** Simulated $BF_{TS}$ time courses for $f_{stim}$ 5, 10 and 20 Hz (**A1**) and 50, 100 and 200 Hz (**A1**) (compare to Fig. 3A).
**B.** Simulated mean $M_{LS,RS} \cdot BF_{LS,RS}$ release time courses for $f_{stim}$ 5, 10 and 20 Hz (**B1**) and 50, 100 and 200 Hz (**B2**) (compare to Fig. 3B).
**C.** Simulated $BF_{LS}$ time courses for $f_{stim}$ 5, 10 and 20 Hz (**C1**) and 50, 100 and 200 Hz (**C1**) (compare to Fig. 3C).
For all simulation shown in **A**–**C**, default model parameters (Table 1) were used.

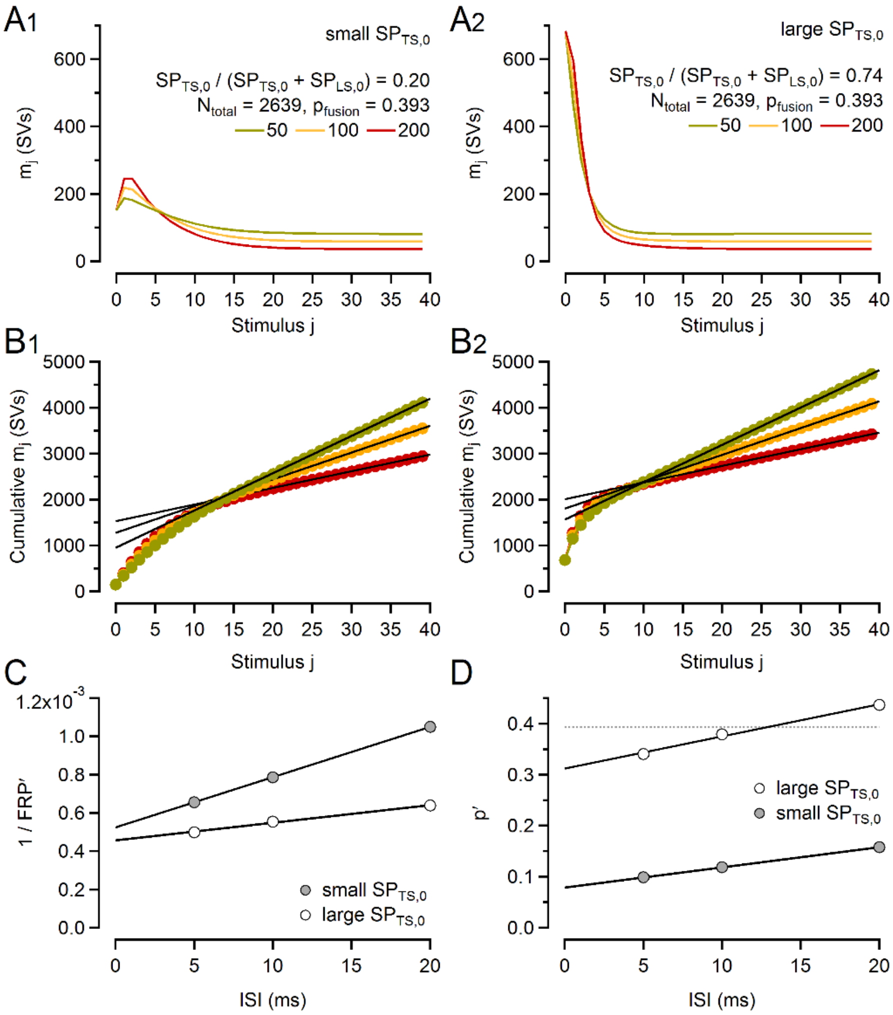


**Fig. S4. 'Traditional' *FRP* and *p* analysis of simulated eEPSC trains for synapses with different $SP_{TS,0}$ fractions misinterprets differences in STP.**

**A.** Time courses of $m_j$ in response to 50, 100 and 200 Hz stimulation simulated using standard values for all model parameters except for $b_2$ which was either increased (**A1**) or decreased (**A2**) such that the fraction $SP_{TS,0}$ / ($SP_{LS,0}$ + $SP_{TS,0}$) was reduced to ~20% or enhanced to ~74%, respectively (replotted from **Fig. 6C**).

**B.** *FRP'* estimates obtained from cumulative $m_j$ for $f_{stim}$ = 50, 100 and 200 Hz for the two simulations shown in **A.**

**C.** Corrected *FRP* estimates derived from back-extrapolations to infinite $f_{stim}$ (ISI = 0 ms) were similar and amounted to 1908 SVs and 2193 SVs for the simulations shown in **A1** and **A2**, respectively. These corrected *FRP* estimates compare favorably to the respective sum $SP_{LS,0}$ + $SP_{TS,0}$ which amounted to 1929 SVs (**A1**) and to 2356 SVs (**A2**).

**D.** Corrected 'traditional' *p* estimates derived from the ratios $m_1$ / *FRP'* differ nearly fourfold for the simulations shown in **A1** ('traditional' $p$ = 0.079) and **A2** ('traditional' $p$ **=** 0.312**)**, despite using the same $p_{fusion}$ value during simulations. For the sequential two-step priming scheme (Fig. 1B), a 'traditional' *p* estimate equals $p_{fusion} \cdot SP_{TS,0}$ / ($SP_{LS,0}$ + $SP_{TS,0}$), corresponding to 0.39 · 0.20 = 0.078 (**A1**) and to 0.39 · 0.74 = 0.290 (**A2**).
