## Supplementary figures and images for "A sequential two-step priming scheme reproduces diversity in synaptic strength and short-term plasticity"

### Figure S1

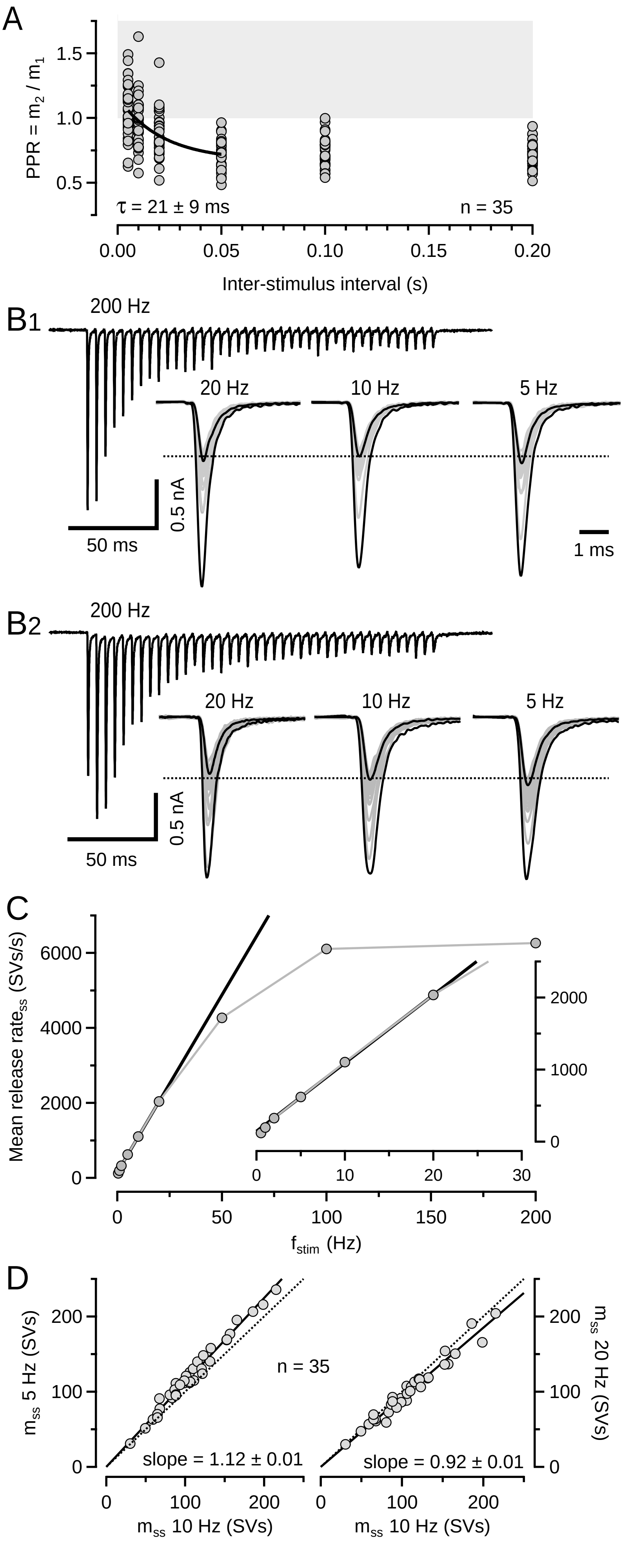


supplemental figure 1

### Figure S2

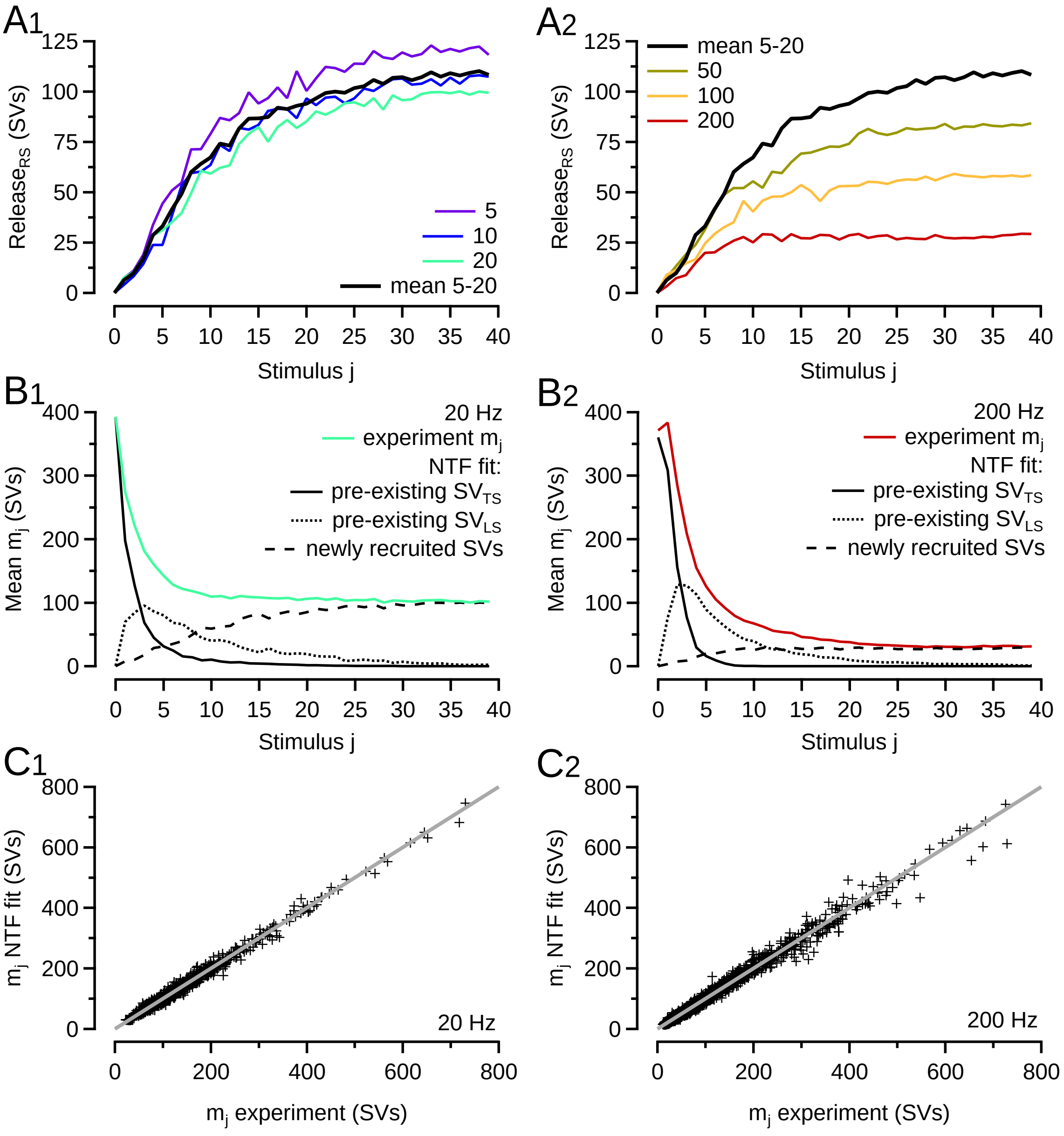


supplemental figure 2

### Figure S3

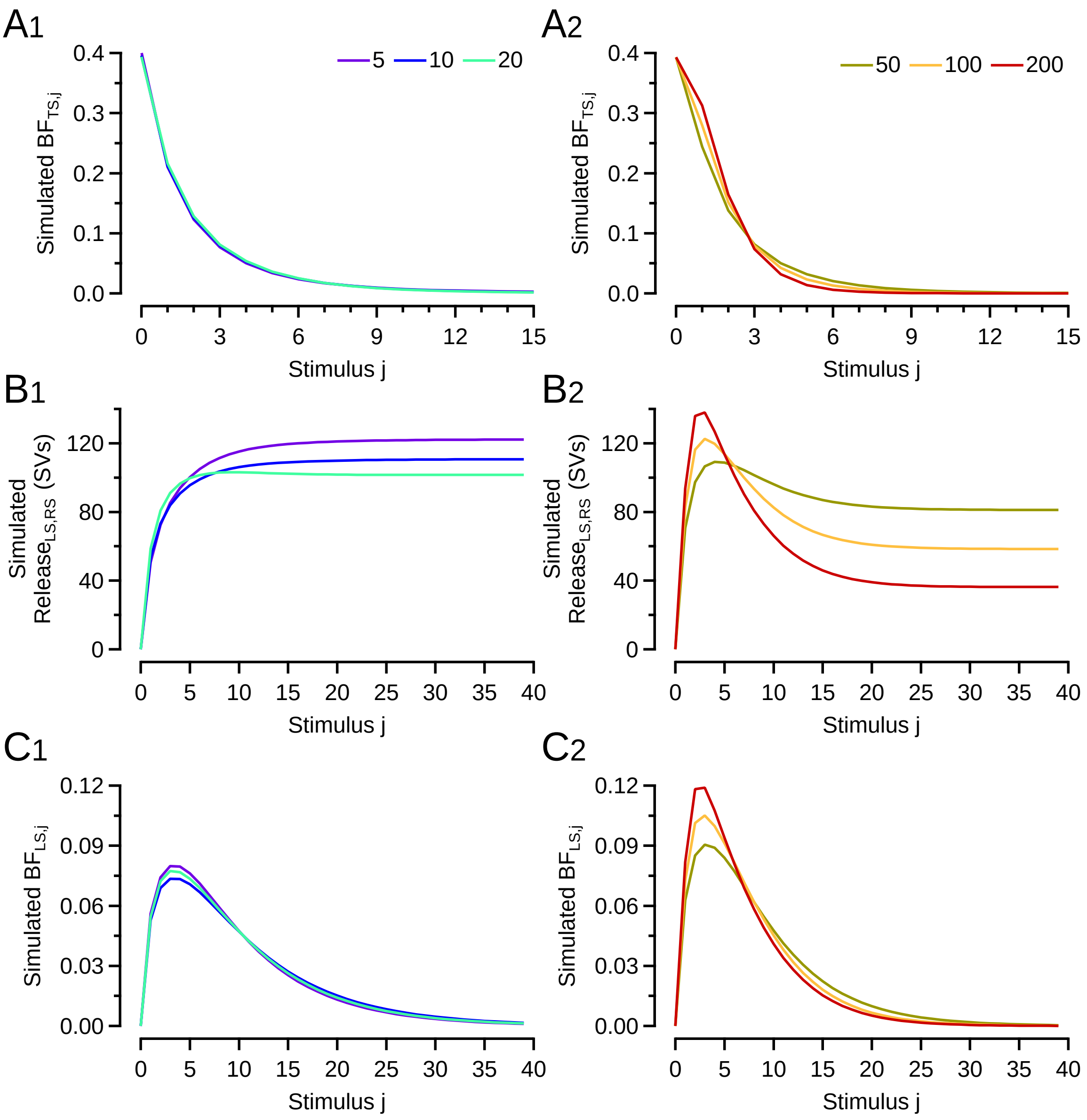


supplemental figure 3

### Figure S4

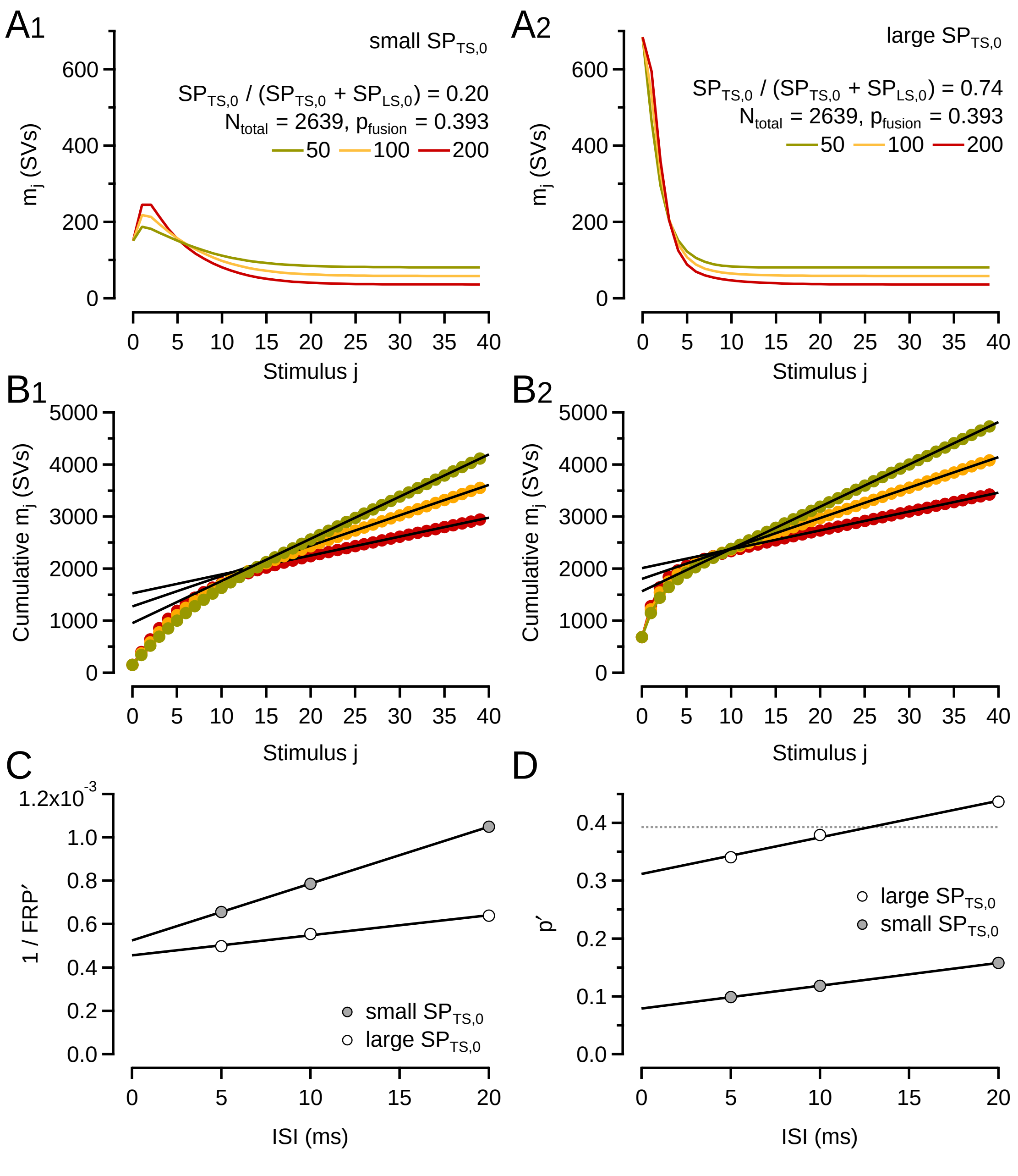


supplemental figure 4
